# Decoupling of transcript and protein concentrations ensures budding yeast histone homeostasis in different nutrient conditions

**DOI:** 10.1101/2023.01.26.525696

**Authors:** Dimitra Chatzitheodoridou, Daniela Bureik, Francesco Padovani, Kalyan Varma Nadimpalli, Kurt M. Schmoller

## Abstract

To maintain protein homeostasis in changing nutrient environments, cells must precisely control the amount of their proteins, despite the accompanying changes in cell growth and biosynthetic capacity. As nutrients are major regulators of cell cycle length and progression, a particular challenge arises for the nutrient-dependent regulation of ‘cell cycle genes’, which are periodically expressed during the cell cycle. One important example are histones, which are needed at a constant histone-to-DNA stoichiometry. Here we show that budding yeast achieves histone homeostasis in different nutrients through a decoupling of transcript and protein abundance. We find that cells downregulate histone transcripts in poor nutrients to avoid toxic histone overexpression, but produce constant amounts of histone proteins through nutrient-specific regulation of translation efficiency. Our findings suggest that this allows cells to balance the need for rapid histone production under fast growth conditions with the tight regulation required to avoid toxic overexpression in poor nutrients.

## Introduction

Protein homeostasis is critical for cell viability and function. Precise regulation of protein amounts must therefore be ensured despite changes in cell morphology, growth, and cell cycle progression that are caused by cellular programs such as cell differentiation, or external factors such as nutrient availability, temperature, oxygen concentration, and osmotic pressure (*1*–*7*).

For example, in nutrient-rich medium, cells typically grow larger and exhibit higher growth rates than in nutrient-poor medium (*6, 8*–*11*). At the same time, rich nutrients lead to elevated ribosome concentrations, which facilitates higher protein synthesis rates and increased biosynthesis (*12, 13*). Nutrients are also major regulators of cell cycle progression and passage through the cell size checkpoints (*14, 15*). Limitation of nutrients or growth factors can induce cell cycle arrest or lengthening of the G1-phase (*1, 16*). For example, budding yeast cells growing slowly on poor medium usually spend more time in G1 until they reach the appropriate size to pass *Start*, the commitment point to cell cycle entry (*17*–*19*). As a result, the durations of S and G2/M-phase become relatively shorter (*20*). In the context of protein homeostasis, such nutrient-induced changes in cell cycle length and progression pose a challenge for the large fraction of ‘cell cycle genes’, which are periodically expressed in a cell-cycle-dependent manner (*21*–*25*). To maintain these proteins at constant concentrations across nutrient conditions, cells would need to adjust protein synthesis rates to changes in cell cycle progression. However, whether and how cells ensure homeostasis of cell-cycle-regulated proteins in changing environments is still unclear.

Histones are a prime example for proteins which are produced in a strongly cell cycle dependent manner and whose concentration needs to be precisely controlled. To couple histone amounts to the genome content and coordinate histone synthesis with DNA replication, the expression of replication-dependent core histone genes is initiated in late G1-phase and then continues throughout S-phase (*26*–*29*). Moreover, in contrast to most proteins, histone production is independent of cell volume, which ensures that histone amounts are tightly coupled to the genome content, even though total protein amounts increase with cell volume (*30*–*33*). In budding yeast, this cell-volume-independent production of histones is already achieved on a transcript level (*31, 32*), at least in part through regulation by the promoter (*31*). This raises the question whether similar regulation also ensures histone homeostasis in changing environments.

Here, we use the budding yeast *Saccharomyces cerevisiae* as a model to understand how cells produce correct amounts of histones in different nutritional environments, despite the accompanying changes in cell size, growth rate and cell cycle fractions (Fig.1A). We find that while cells maintain constant stoichiometry between DNA and histone proteins, histone transcript concentrations are paradoxically decreased in poor compared to rich nutrients – potentially to avoid histone overexpression, which is more toxic in poor growth conditions. This decoupling of histone mRNA and protein amounts across different nutrients implies the requirement for nutrient-dependent regulation of histone translation efficiency. By combining single-cell approaches with population-level analysis, we were able to show that histone promoters are sufficient to mediate nutrient-dependent transcript regulation, and its compensation by translation.

## Results

### Histone protein concentrations decrease with cell volume across nutrient conditions

To determine the impact of nutrient availability on the regulation of histone protein concentrations, we grew wild-type haploid cells on synthetic complete (SC), yeast peptone (YP) and minimal medium (SD), containing glucose (D) or galactose (Gal) as fermentable and glycerol and ethanol (GE) as non-fermentable carbon sources. The selected set of seven different growth media resulted in a wide variety of growth rates, cell sizes and cell cycle phase distributions. Population doubling times ranged from 1.3 h to 6.6 h, with cells growing fastest on YPD and slowest on non-fermentable minimal growth medium (SDGE) (Fig. 1B). The mean cell volume was largest in YPD at about 62 fL and smallest in SCGE at 46 fL (Fig. 1C). To determine the fraction of cells in different cell cycle phases, we quantified the DNA content with flow cytometry. The percentage of cells in G1-phase notably increased in media with non-fermentable carbon sources, leading to smaller fractions of cells in S- and G2/M-phases (Fig. 1D). Using the population doubling times to estimate absolute durations from the cell cycle fractions indicates S-phase duration varies between 29 min to 88 min (Supplementary Fig. 1A).

**Figure 1.**
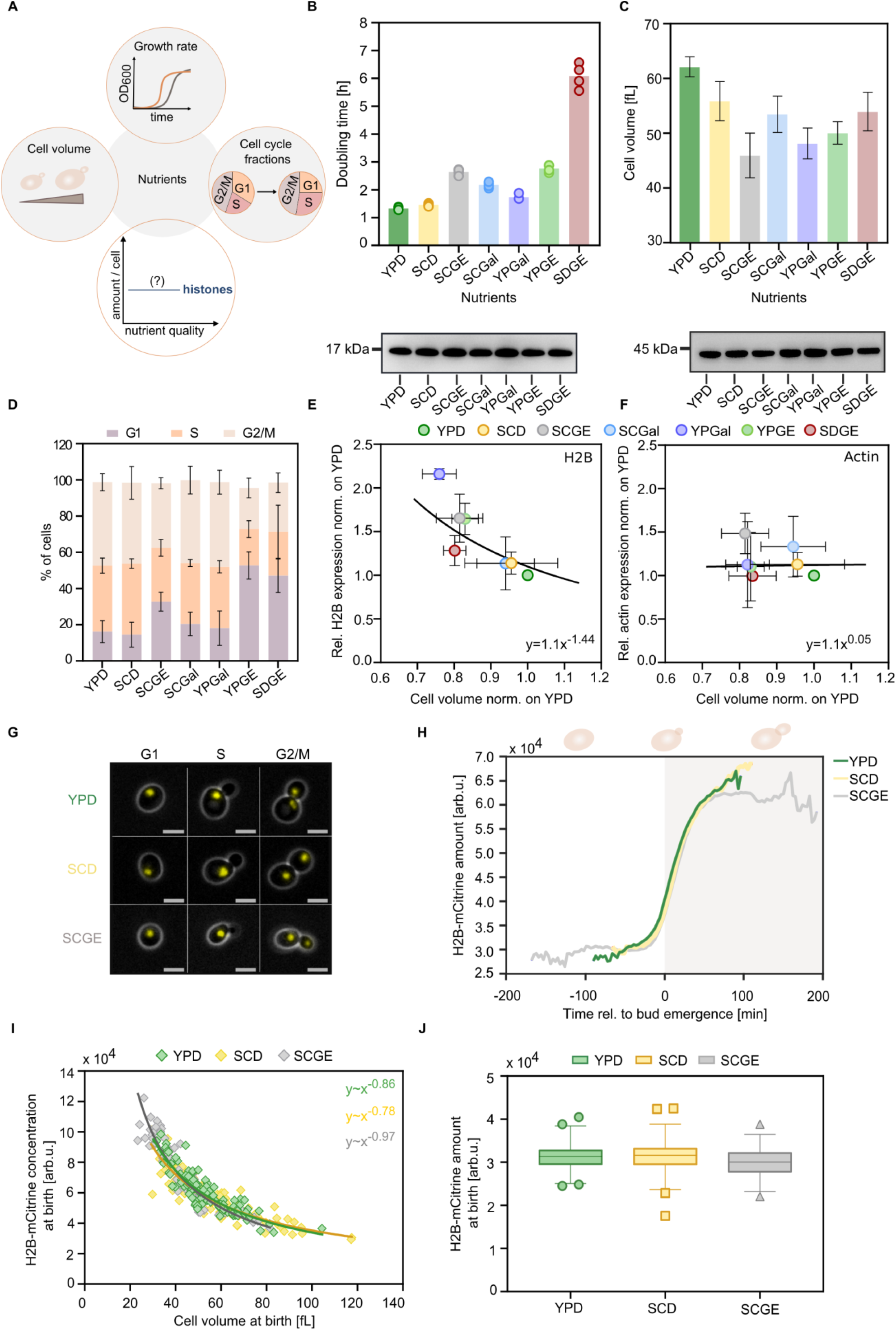
Single cell and population level-analyses reveal that histone protein concentrations decrease with cell volume across nutrient conditions, despite the nutrient-induced changes in cell growth. (**A**) The nutrient environment has a pronounced influence on various aspects of cell growth, such as cell size, growth rate and cell cycle progression. As histones are cell-cycle-regulated proteins, the question arises of whether cells maintain histone homeostasis despite the nutrient-induced changes in cell growth. To determine the impact of different nutrients on the regulation of histone proteins, we grew cells on synthetic complete (SC), yeast peptone (YP) and minimal medium (SD), containing glucose (D) or galactose (Gal) as fermentable and glycerol and ethanol (GE) as non-fermentable carbon sources. (**B**) Doubling times calculated from growth curves of exponentially growing cell populations in different growth media. The bar graphs represent the mean of n = 4 independent replicates, each shown as an individual dot. (**C**) Mean cell volumes of cells growing in different nutrients, measured with a Coulter counter. Bar plots with error bars indicate mean values and standard deviations across n = 6 replicate measurements. (**D**) Flow cytometry analysis of the nutrient-dependent cell cycle distributions (percentage of cells in G1-, S- and G2/M-phase) based on quantification of the DNA content. Error bars represent the standard deviation of n = 5 independent measurements. (**E-F**) Protein bands of (**E**) histone H2B and (**F**) actin protein were quantified by western blot in different growth media and normalized to total proteins as determined from Ponceau stains. For each condition, total protein content was extracted from equal number of cells. The relative protein expression is shown as a function of the relative nutrient-specific cell volume. Line shows fit with equation parameters obtained from linear regression on the double-logarithmic data. (**G**) Representative live cell fluorescence and phase-contrast images of G1, S and G2/M cells with *mCitrine*-tagged H2B (both Htb1 and Htb2 tagged), growing in three nutrient conditions. The scale bars represent 5 μm. (**H**) Mean amounts of H2B-mCitrine during the first cell cycle of newborn cells in YPD (n_YPD_ = 87), SCD (n_SCD_ = 83) and SCGE (n_SCGE_ = 55). Single cell traces are aligned at bud emergence (t = 0). (**I**) H2B-mCitrine concentrations at birth are plotted as a function of cell volume for three nutrient conditions. Line shows fit with equation parameters derived from a linear regression on the double-logarithmic data. (**J**) H2B-mCitrine amounts at birth in the different growth media. Box plots represent median and 25th and 75th percentiles, whiskers indicate the 2.5th and 97.5th percentiles and symbols show outliers.

To test if cells maintain a constant histone-to-DNA stoichiometry in the different nutrients, we measured the histone H2B protein levels by western blot analysis. For all media, we extracted total protein content from equal number of cells and used Ponceau S staining for quantification (Supplementary Fig.1B). Consistent with previous studies (*34, 35*), we observe a decrease in total protein abundance in nutrient-poor conditions (Supplementary Fig.1C). To compare the relative H2B protein expression in all growth media, we then normalized the histone protein amounts to total protein amounts. Across the different nutrients, we find that the histone H2B protein concentration decreases in inverse proportion with cell volume (Fig. 1E), indicating that histone protein synthesis is coordinated with genomic DNA content rather than cell volume. In contrast, actin is maintained constant at a cell-volume independent concentration (Fig. 1F). This is consistent with the fact that actin expression scales with cell volume to ensure constant concentrations during growth (*31, 32*).

### In different growth conditions, histone protein concentrations decrease with cell volume on a single cell level

The above data shows histone protein concentrations decrease with cell volume to maintain constant amounts across nutrient conditions. To confirm that this is also true within a given cell cycle stage, we performed microfluidics based live cell microscopy and monitored the synthesis of histone H2B over time in individual haploid cells. For this purpose, we endogenously tagged *HTB1* and *HTB2*, the two genes encoding for the core histone H2B, with the fluorescent protein mCitrine. We then measured cell volume and total mCitrine intensity in new born daughter cells by timelapse microscopy across three nutrient conditions (Fig. 1G, H).

In all growth media, H2B-mCitrine amounts per cell are constant during early G1 and increase approximately two-fold during S-phase before they reach a plateau in G2M. As expected, the cell cycle duration strongly depends on the nutrient condition, with cells in SCGE exhibiting the longest cell cycle (Fig. 1H). Yet for all conditions, H2B protein concentration at birth decreases with cell volume because equal total histone amounts are produced in the mother cells (Fig. 1I, J). This was also confirmed by flow cytometry measurements of total H2B-mCitrine amounts in G1 in all seven growth media (Supplementary Fig. 2). Thus, our experiments with fluorescently tagged H2B support the conclusion that histone protein amounts are maintained constant across nutrient conditions. However, we note that while the fluorescent tagging of both histone genes did not noticeably affect population doubling times, we detected increased mean cell volumes and elevated *HTB1* and *HTB2* transcript concentrations (Supplementary Fig. 3A, B, C). Thus, the exact mode of histone regulation in these strains has to be interpreted with care.

**Figure 2.**
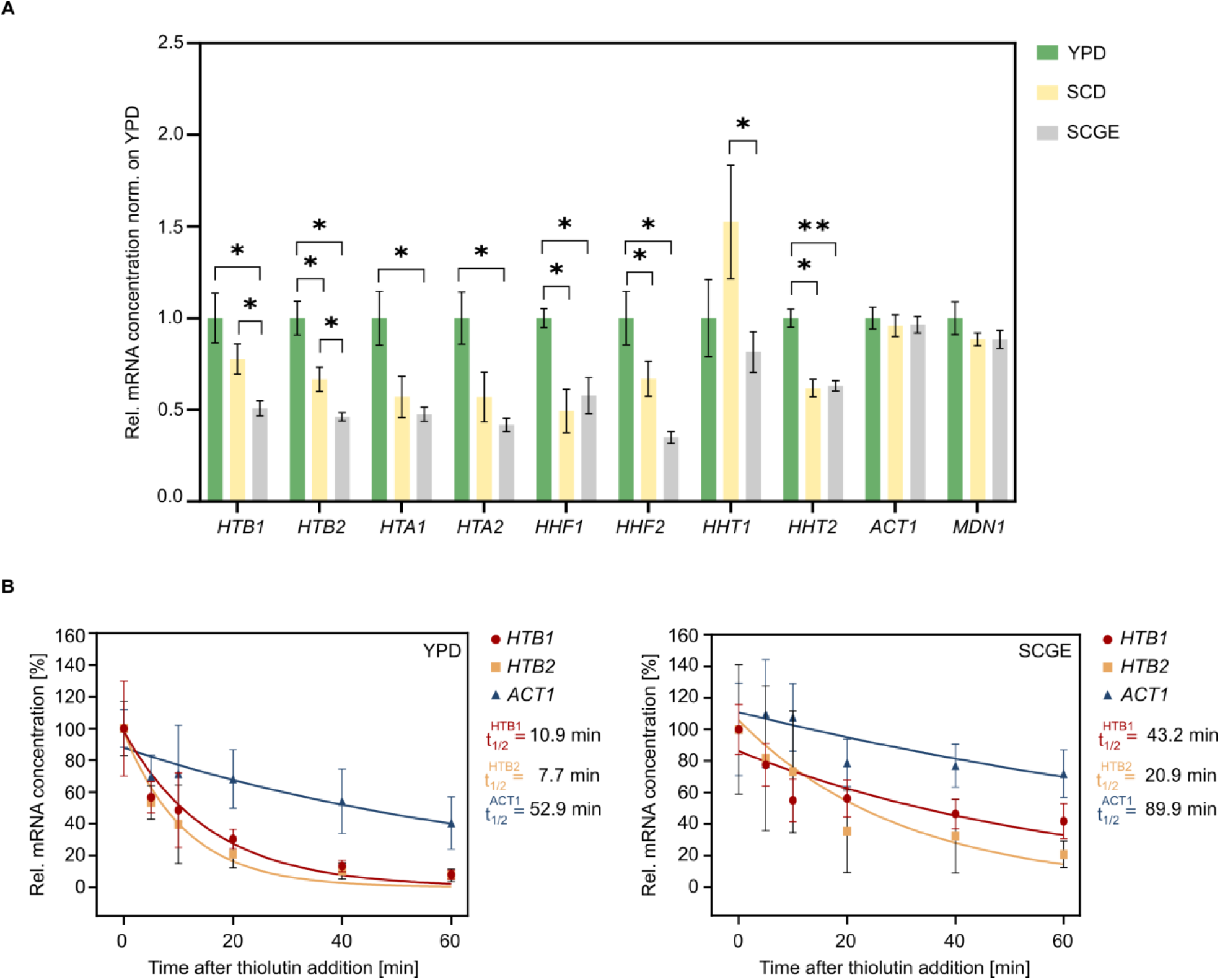
Histone transcription is downregulated in poor nutrients. (**A**) RT-qPCR was used to quantify the mRNA concentrations of the core histone genes and the control genes *ACT1* and *MDN1* in different nutrient conditions. mRNA concentrations were normalized on *RDN18* and are shown as fold changes compared to YPD. Bars represent the mean values of at least 4 independent biological replicates; error bars indicate standard errors. Significances were determined by an unpaired, two-tailed t-test for datasets that follow a Gaussian distribution or a Mann-Whitney test for datasets that are not normally distributed. (* p<0.05, ** p<0.01). (**B**) Transcription inhibition experiments suggest that histone mRNA stability increases in poor nutrient conditions. The mRNA half-lives of *HTB1, HTB2* and *ACT1* were determined by adding the RNA polymerase inhibitor thiolutin to cells growing in different growth media and then measuring mRNA concentrations (normalized on *RDN18*) over time by RT-qPCR. Relative mRNA concentrations were normalized on the initial concentration at time=0. For each time point, the mean and standard deviation of at least 4 biological replicates is plotted. Lines show single exponential fits to the individual data points of all replicates.

**Figure 3.**
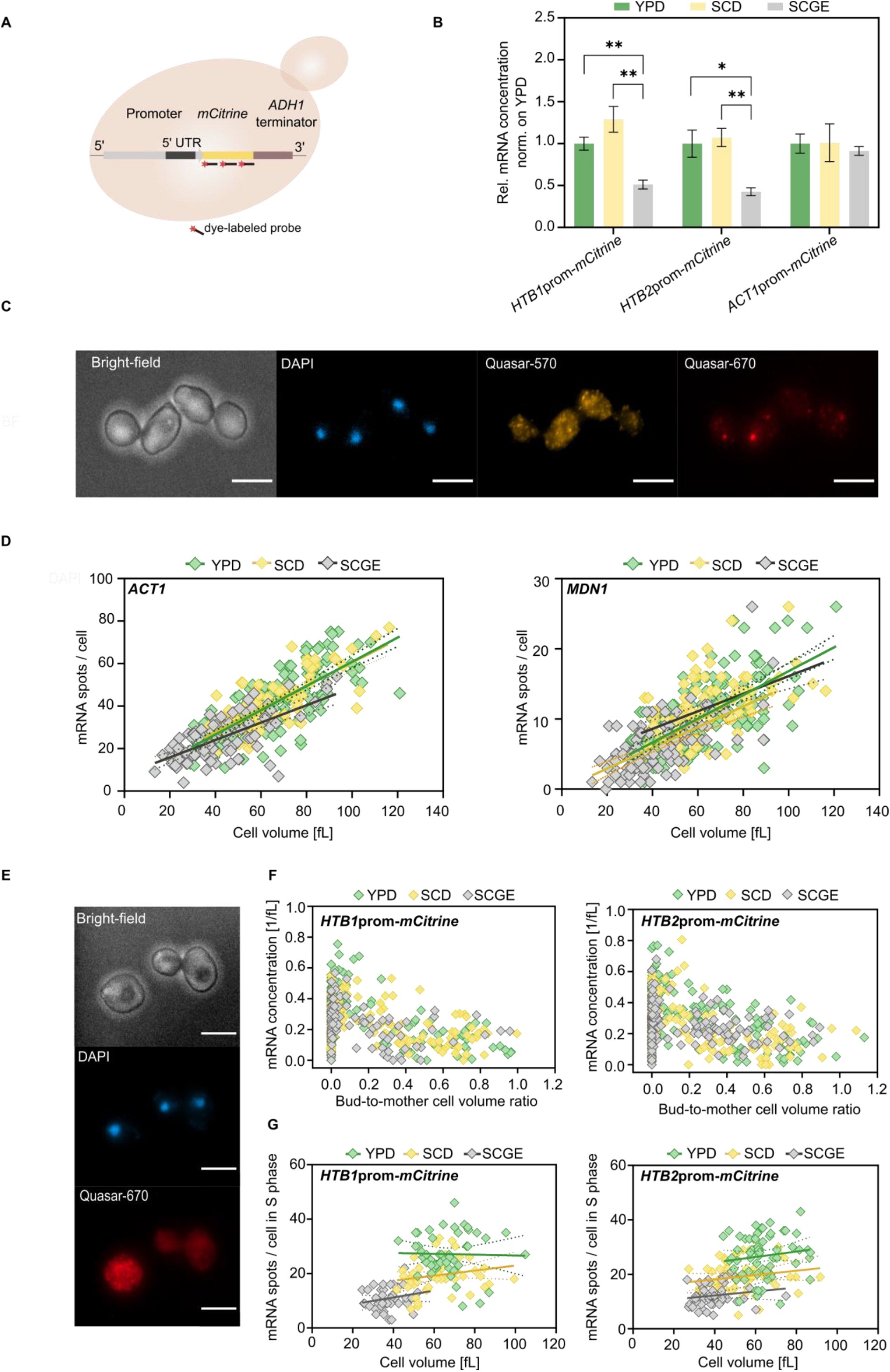
Transcript amounts expressed from histone promoters are independent of cell volume in all nutrient conditions. (**A**) An *mCitrine* reporter gene driven by the promoter of interest (including the 5’ UTR) and regulated by the *ADH1* terminator was endogenously integrated in the *URA3* locus of wild-type haploid cells. smFISH was performed to study cell-volume- and cell-cycle-dependent gene expression in different nutrient conditions on a single cell level (Supplementary Fig. 6). Schematic representation of fluorescently labeled probes binding to the target mRNA. (**B**) Relative mRNA concentrations of *mCitrine* expressed from the *HTB1, HTB2* or *ACT1* promoter as determined by RT-qPCR. Bars represent mean fold changes with respect to YPD and error bars indicate standard errors of at least 4 replicate measurements. Statistical significance was calculated by an unpaired, two-tailed t-test (* p<0.05, ** p<0.01). (**C**) Epifluorescence microscopy was performed to detect *ACT1* and *MDN1* transcripts targeted with Quasar- 570- (yellow) and Quasar-670-labelled (red) probes, respectively. Representative images of cells grown in YPD are shown. Cell nuclei were stained with DAPI (blue) and bright-field images were used to estimate cell volume. The scale bars represent 5 μm. (**D**) Nutrient-dependent mRNA amounts of *ACT1* and *MDN1* per cell (n_YPD_ = 176, n_SCD_ = 87, n_SCGE_ = 98) as a function of cell volume. Lines show linear fits; dashed lines indicate the 95% confidence intervals. (**E**) Representative images of cells expressing *HTB1*prom-*mCitrine* in YPD. Transcripts were detected using probes labeled with Quasar-670 (red). Nuclear DNA was stained with DAPI (blue). The scale bars represent 5. (**F**) mRNA concentration of *mCitrine* expressed from an additional *HTB1* (n_YPD_ = 149, n_SCD_ = 158, n_SCGE_ = 95) or *HTB2* promoter (n_YPD_ = 161, n_SCD_ = 194, n_SCGE_ = 170) plotted against the corresponding bud-to-mother cell volume ratio in different nutrients. (**G**) Number of *mCitrine* mRNA spots per cell (*HTB1*prom*-mCitrine*; n_YPD_ = 49, n_SCD_ = 51, n_SCGE_ = 39), (*HTB2*prom- *mCitrine*; n_YPD_ = 64, n_SCD_ = 59, n_SCGE_ = 50) as a function of cell volume during S-phase. Here, cells with one nucleus and a bud-to-mother volume ratio < 0.3 were considered to be in S-phase. Lines show linear fits; dashed lines indicate the 95% confidence intervals.

### Histone transcript concentrations are downregulated in poor nutrient conditions

Our results demonstrate that cells couple histone protein amounts to the genome content despite the nutrient-induced changes in cell growth and cell cycle progression. Since previous studies showed that the cell-volume-independent coupling of histone amounts to DNA content is ensured at the transcript level (*31, 32*), we asked whether the nutrient-dependent histone homeostasis observed here is also achieved at the transcript level. In that case, we would predict that histone transcript amounts would be constant across nutrient conditions, resulting in an increased mRNA concentration for cells cultured in nutrient-poor conditions according to their relatively smaller cell size. To test this, we performed RT-qPCR and quantified histone transcript concentrations relative to total RNA in three nutrient conditions (Fig. 2A). In contrast to our expectations, we found that histone transcript concentrations are significantly decreased in poor compared to rich growth media. At the same time, cells maintain constant mRNA concentrations of the control genes *ACT1* and *MDN1*.

Our findings indicate that while the amount of histone proteins is tightly coupled to the DNA content, histone mRNA expression shows an unexpected dependence on the nutrient conditions. The apparent decoupling of histone protein and mRNA abundance suggests that medium-specific regulation of protein translation or stability is required to ultimately ensure histone homeostasis in different nutritional environments.

### Transcription accounts for nutrient-dependent downregulation of histone transcripts in nutrient-poor conditions

As mRNA concentration is set by transcription and mRNA degradation, we sought to determine the contribution of these opposite reactions to the observed nutrient-dependent regulation of histone mRNA levels. More precisely, we studied histone and *ACT1* mRNA stability in rich and poor medium by inhibiting global transcription with thiolutin and monitoring the remaining mRNA by RT-qPCR over time. We then fitted a single exponential function to the mRNA decay curves to determine the mRNA half-lives (Fig. 2B). Consistent with previous studies (*36, 37*), we found that histone mRNAs in YPD have short half-lives, whereas *ACT1* mRNA is more stable. Moreover, we showed that both histone and *ACT1* mRNAs are significantly more stable in poor compared to rich nutrient conditions. Again, this is consistent with previous studies that showed that transcription and degradation rates of many genes tend to increase in fast growth conditions, ensuring constant mRNA concentrations (*38, 39*).

Taken together, our results suggest that while the nutrient environment affects both histone mRNA synthesis and degradation, it is histone transcription that accounts for the nutrient-dependent downregulation of histone transcripts in nutrient-poor conditions.

### Histone promoter determines nutrient-dependence of histone transcript concentrations

Previously, we have shown that histone promoters can mediate the coordination of histone transcripts with genomic DNA content despite changes of cell volume (*31*). We therefore asked whether histone promoters are also sufficient for the nutrient-dependent regulation of histone transcript concentrations. To test this, we created strains with an endogenously integrated *mCitrine* reporter driven by an additional *HTB1, HTB2* or *ACT1* promoter (including the 5’ UTRs), respectively (Fig. 3A). Indeed, we found that the reporter mRNA concentrations decrease in nutrient poor conditions, similar to the endogenous histone concentrations. In contrast, if expressed from the *ACT1* promoter, the *mCitrine* transcripts are kept at constant, nutrient-independent concentrations (Fig. 3B).

### Histone transcripts amounts are independent of cell volume in each different nutrient condition

So far, we have established that histone promoters can be sufficient to mediate nutrient-dependent regulation, but it is unclear whether the histone mRNA synthesis is uncoupled from cell volume for cells grown on each medium. To test this, we performed single-molecule fluorescence in situ hybridization (smFISH) combined with bright-field fluorescence microscopy, which allows studying cell-volume- and cell-cycle-dependent gene expression in different nutrient conditions on a single cell level (Fig. 3A). We used bright-field images to determine cell volume and assigned each cell to a cell cycle stage based on the calculated bud-to-mother volume ratio, as well as the morphology and number of DAPI-stained nuclei. First, we quantified the nutrient-dependent mRNA concentrations of *ACT1* and *MDN1*, two representatives of scaling gene expression (*31, 32*) (Fig. 3C, Supplementary Fig. 4A). As anticipated, both genes were continuously expressed throughout the cell cycle at constant concentrations (Supplementary Fig. 4B, C), and transcript amounts were increasing proportionally to cell volume (Fig. 3D). Moreover, we found similar mRNA copy numbers at a given cell volume regardless of the nutrient condition. This highlights the importance of maintaining appropriate concentrations despite the nutrient-induced changes in cell growth.

**Figure 4.**
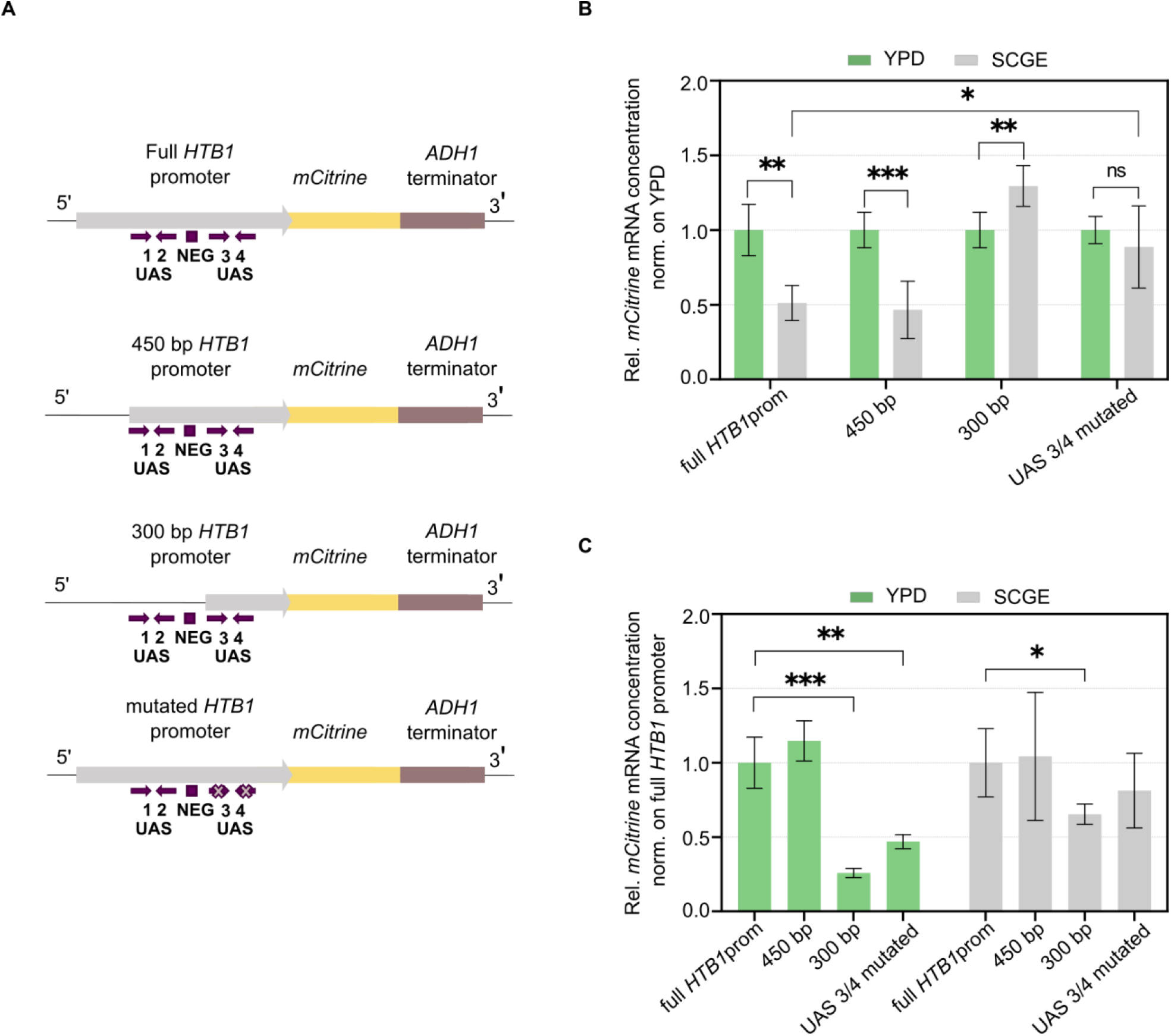
Spt10 binding sites in the *HTB1* promoter are required for the nutrient and cell-size-dependence of transcript concentrations. (**A**) Schematic representation of mCitrine expressed from the full *HTB1* promoter, the 450 bp and 300 bp truncations as well as the *HTB1* promoter with mutated Spt10 binding sites in UAS3 and UAS4. The arrows indicate the location and orientation of the UAS elements and the boxes show the NEG region. (**B**) RT-qPCR analysis of the *mCitrine* mRNA concentrations (normalized on *RDN18*) in different growth media. The results are shown as fold changes with respect to YPD in the respective media. Bar plots with error bars indicate mean values and standard deviations across n = 5-8 independent replicates. (**C**) Quantification of *mCitrine* mRNA concentrations (normalized on *RDN18*) by RT-qPCR. The results are shown as fold changes compared to the full *HTB1*prom-*mCitrine*. Bar plots with error bars indicate mean values and standard deviations across n = 5-8 independent replicates. Significances were determined by an unpaired, two-tailed t-test for datasets that follow a Gaussian distribution or a Mann-Whitney test for datasets that are not normally distributed. (* p<0.05, ** p<0.01).

In contrast to *ACT1* and *MDN1*, histone biogenesis is regulated in a cell cycle-dependent manner, as it is tightly coupled to DNA replication. Consequently, histone genes are transcriptionally active in late G1 and S-phase (*26, 40, 41*). Previous studies have shown that histone promoters are sufficient to mediate periodic transcription, resulting in an mRNA expression peak during early-to-mid S-phase, which is coordinated with genome content rather than cell volume (*31, 42, 43*). Using smFISH, we quantified the nutrient-dependent mRNA concentrations of *mCitrine* expressed from the *HTB1* or *HTB2* promoter (Fig. 3E, Supplementary Fig. 4D, E). In all growth conditions, we observe increased *mCitrine* mRNA concentrations in S-phase, before they eventually decline as cells progress further through the cell cycle (Fig. 3F). We also find that for each nutrient condition, the mRNA peak-expression during S-phase is uncoupled from cell volume, leading to constant transcript amounts per cell. Yet, in nutrient-poor conditions, the mRNA copy number at a given cell volume is significantly lower than in rich conditions (Fig. 3G). This nutrient-dependent decrease of mRNA abundance in S-phase in poor nutrients is consistent with our RT-qPCR-based results quantifying average mRNA concentration in asynchronous populations. Further confirming the distinct nutrient-dependence mediated by histone promoters, we found that if expressed from an *ACT1* promoter, *mCitrine* transcript amounts per cell increase with cell volume in all three conditions, but are independent of the nutrient condition – similar to the transcripts of endogenous *ACT1* (Supplementary Fig. 5).

**Figure 5.**
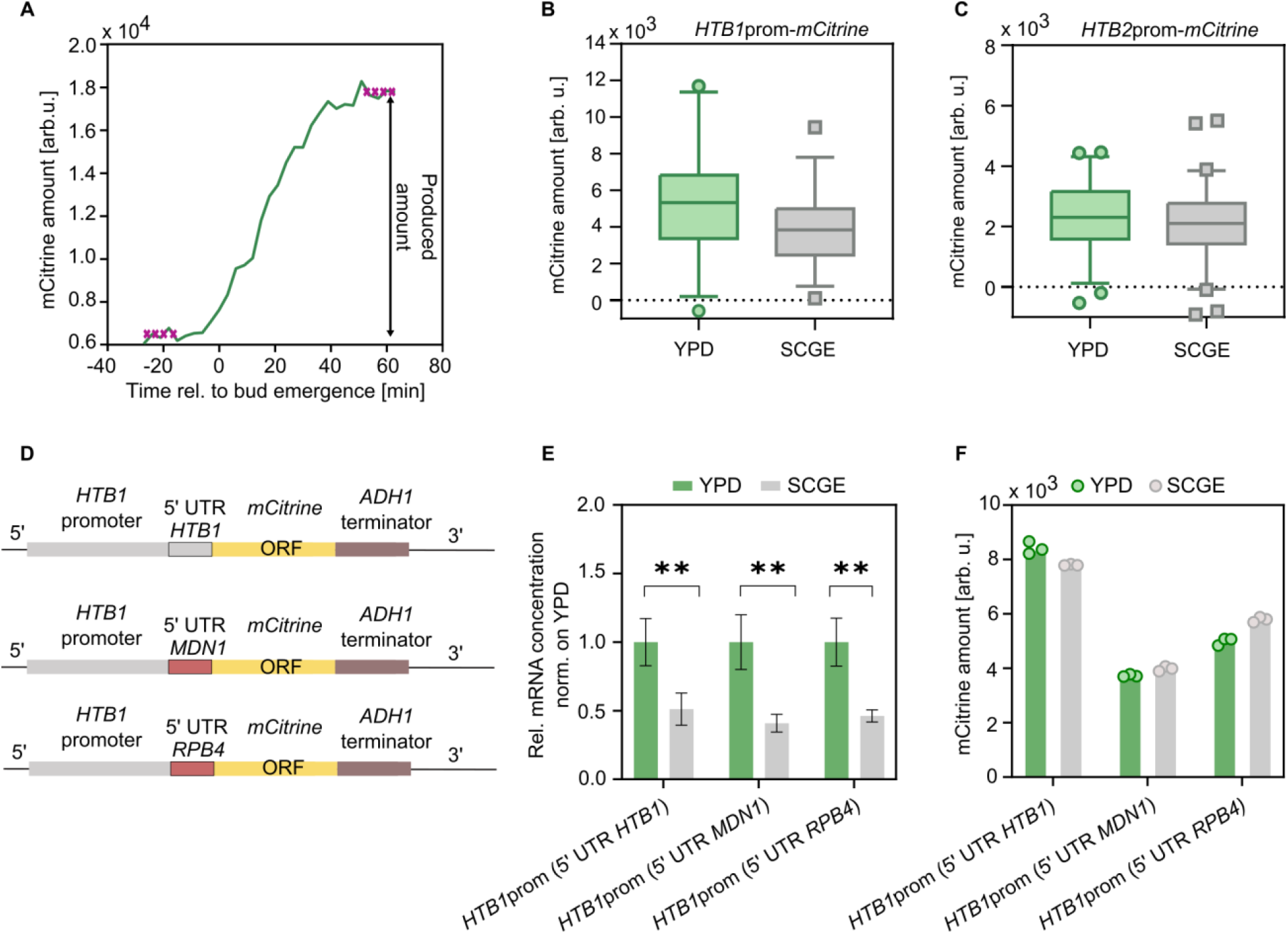
mCitrine protein expression driven by H2B promoters is uncoupled from nutrient-dependent mRNA levels. (**A**) Protein amounts of mCitrine expressed from the *HTB1* and *HTB2* promoter during the cell cycle were measured in rich and poor growth medium using live-cell fluorescence microscopy. Representative fluorescence intensity trace of a haploid cell expressing *HTB1*prom-*mCitrine* in YPD. The mCitrine amounts produced during the cell cycle were calculated as the difference between the median of the first four time points and the median of the last four time points. (**B-C**) Protein amounts of mCitrine expressed from the *HTB1* and *HTB2* promoter were measured using live-cell fluorescence microscopy. Box and whisker plots represent the distribution of total mCitrine amounts produced during the cell cycle, under different nutrient conditions. Box plots represent median and 25th and 75th percentiles, whiskers indicate the 2.5th and 97.5th percentiles and symbols show outliers. (**D**) To examine the influence of the *HTB1* 5’ UTR on the nutrient-dependent mCitrine expression from the *HTB1* promoter, the 5’ UTR was replaced with the 5’ UTRs of *MDN1* and *RPB4*, respectively. (**E**) RT-qPCR analysis of the *mCitrine* mRNA concentrations (normalized on *RDN18*) in different growth media. Results are shown as fold changes compared to YPD. Bar plots with error bars indicate mean values and standard deviations of at least 4 independent replicates (** p<0.01). (**F**) mCitrine protein amounts as determined by flow cytometry. Bars represent mean fluorescence intensities of 3 biological replicates, each depicted as individual dot.

### Histone promoter truncation changes the nutrient-dependence of transcript concentrations

In *Saccharomyces cerevisiae*, the expression of core histones is controlled by positive and negative regulatory elements in the promoter regions. Positive transcriptional regulation is mediated by transcription factors including Spt10 and SBF that bind to the upstream activating sequences (UASs) and activate transcription periodically. In contrast, the histone regulatory (HIR) complex acts as a negative transcriptional regulator that represses histone gene expression by binding to the NEG element, present in all core histone promoters except the *HTA2–HTB2* promoter pair (*41, 44*).

Previously, we found that decreasing promoter strength can alter the cell volume-dependence of histone promoters, resulting in a promoter-mediated scaling of gene expression with cell volume (*31*). Specifically, a truncated *HTB1* promoter, consisting only of 300 bp and lacking part of the UASs and the NEG element, drives the expression of mCitrine in a cell-volume dependent manner. Motivated by these findings, we sought to determine whether different truncations of the *HTB1* promoter also induce changes in the nutrient-dependence of the *mCitrine* mRNA levels. For this purpose, we used haploid strains expressing mCitrine driven by a 450 and 300 bp *HTB1* promoter (including the 5’ UTR), respectively, each truncated from the 5′-end (*31*) (Fig. 4A).

Our analysis revealed that, similar to the full-length promoter, the 450 bp truncation showed reduced *mCitrine* mRNA concentrations in poor compared to rich condition (Fig. 4B). In contrast, for the 300 bp truncation we found that *mCitrine* mRNA concentrations considerably decreased in rich growth medium (Fig. 4C), becoming significantly lower than in poor medium (Fig. 4B). All strains tested in this experiment exhibited similar cell volumes and doubling times in the respective growth media (Supplementary Fig. 7A, B). Our results therefore imply that the loss of the 150 bp sequence between the 450 bp and 300 bp truncation of the *HTB1* promoter leads to a marked change in the nutrient-dependence of the *mCitrine* transcripts. Given that this 150 bp sequence includes the UAS1 and UAS2 as well as the NEG element, it is possible that those regulatory elements contribute to the nutrient-dependent regulation of histone transcripts.

**Figure 6.**
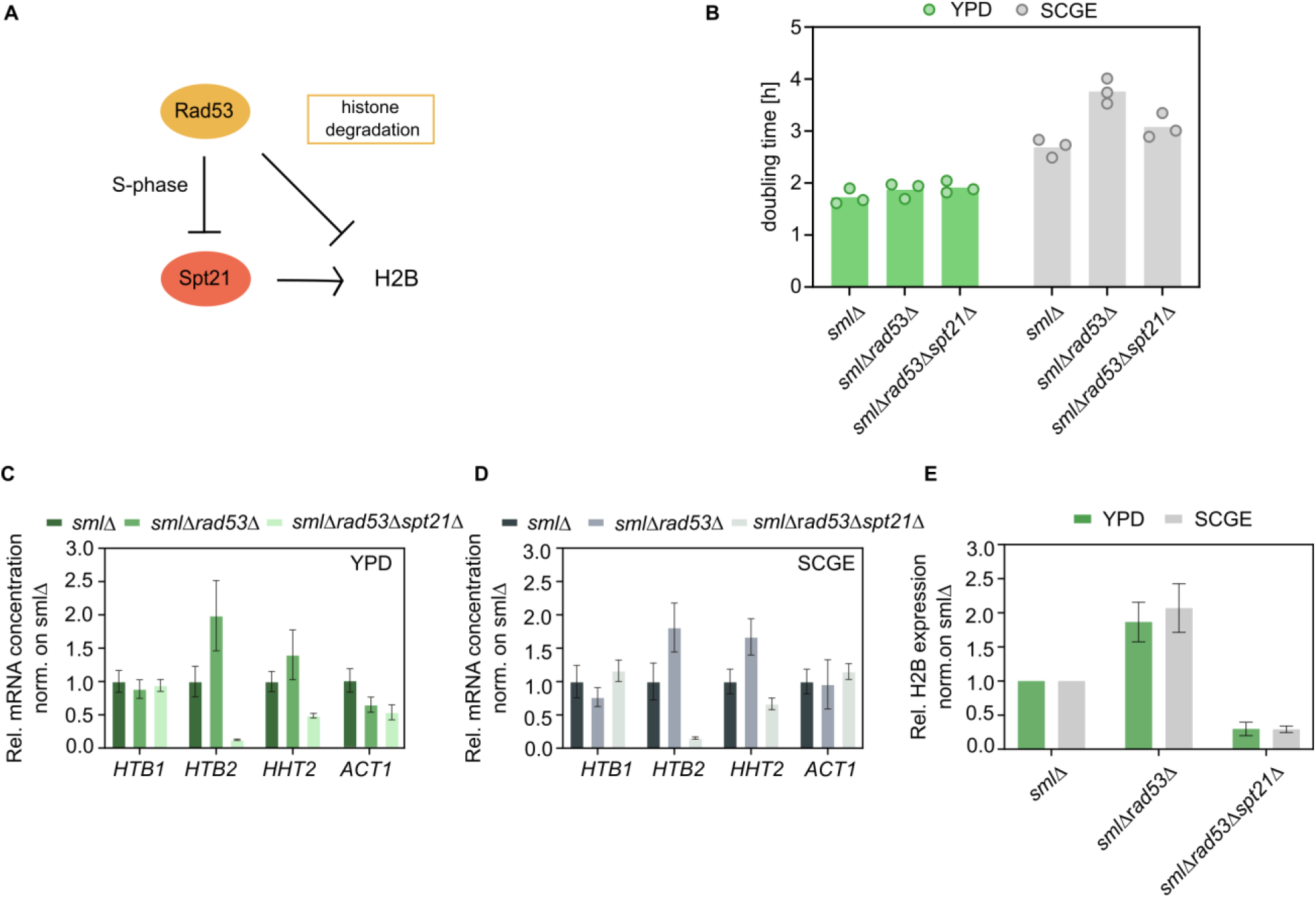
Cells growing on non-fermentable carbon sources are more sensitive to histone overexpression than in nutrient-rich conditions. (**A**) Rad53 is required for the degradation of excess histones and also regulates histone levels by inhibiting the transcription activator Spt21. (**B**) Doubling times calculated from growth curves of *sml1Δ, sml1Δrad53Δ* and *sml1Δrad53Δspt21Δ* cells growing on fermentable and non-fermentable carbon sources. Bars represent the mean of n = 3 independent measurements shown as individual dots. (**C-D**) mRNA concentrations of *HTB1, HTB2, ACT1* and *HHT2* (normalized on *RDN18*) for *sml1Δrad53Δ* and *sml1Δrad53Δspt21Δ* cells are shown as fold changes with respect to *sml1Δ*, in YPD (**C**) and SCGE (**D**). Bar plots with error bars indicate mean values and standard errors across n = 3-5 biological replicates. (**E**) H2B protein levels, normalized on total protein, as determined by western blot analysis. Bars represent mean fold changes with respect to *sml1Δ* and error bars indicate standard errors of at least 4 replicates.

**Figure 7.**
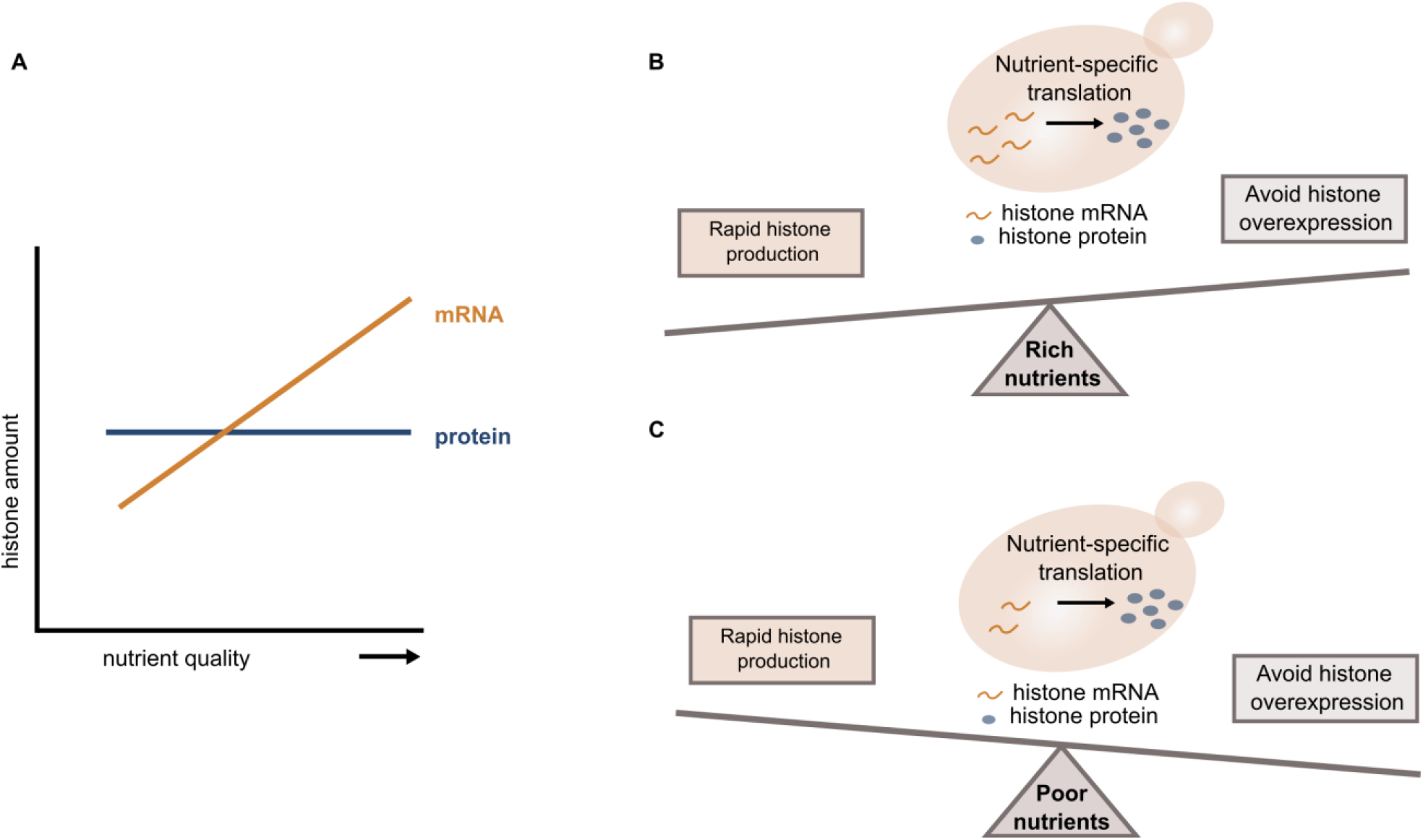
Histone homeostasis in different nutrient environments. (**A**) While cells maintain constant amounts of histone proteins across changing environments, the histone mRNA expression is increased in nutrient-rich conditions. (**B-C**) Decoupling of protein and mRNA abundance is achieved through nutrient-dependent translation, allowing cells to balance the need for rapid histone production in rich growth media (**B**) with the tight regulation required to prevent histone overexpression in poor growth media (**C**).

### Spt10 is required for the nutrient and cell-size-dependence of transcript concentrations

We next asked whether the transcription factor Spt10, which specifically binds to the UASs, is required for the increase of histone mRNA levels in rich growth media. To test this, we constructed a haploid strain carrying an additional copy of the *HTB1* promoter driving mCitrine, in which we mutated the Spt10 binding sites of the UAS3 and UAS4 elements (*43*) (Fig. 4A). We then quantified the *mCitrine* mRNA concentrations by RT-qPCR and found that mutation of the two UASs altered the nutrient-dependence, leading to similar transcript concentrations in rich and poor growth medium (Fig. 4B). These results suggest a role of Spt10 in regulating histone gene expression in response to the nutrient environment.

### Histone promoter is sufficient to compensate for nutrient-dependent histone transcript regulation

We have shown that despite the nutrient-dependence of histone transcript concentrations, cells maintain a constant histone-to-DNA stoichiometry across changing environments. This suggests that additional regulation of translation efficiency or protein stability is required to achieve constant histone amounts across different nutrient conditions. We next asked whether histone promoters can also mediate this regulation at the protein level. We tested this by performing microfluidics based live cell microscopy and measuring the protein amounts of mCitrine expressed from the *HTB1* and *HTB2* promoter, respectively. In contrast to the endogenous histones, which are evenly distributed between the mother and daughter cell during division, mCitrine is partitioned along with the cytoplasm, in proportion to cell volume (*32*). Due to the asymmetric division of budding yeast, differently sized G1 cells could therefore inherit different amounts of mCitrine. Consequently, we refrained from comparing mCitrine amounts at birth and instead calculated the amounts produced during the cell cycle in rich and poor growth medium (Fig. 5A). Our results reveal that while the mRNA amounts of *mCitrine* at a given cell volume are approximately two-fold lower in poor compared to rich medium (Fig. 3G), the produced protein amounts are much more similar between conditions (Fig. 5B, C). Thus, consistent with the experimental findings for the endogenous histones, the nutrient-dependence of mCitrine proteins expressed from H2B promoters is uncoupled from the mRNA levels. The fact that this regulation occurs also for the mCitrine reporter suggests that it is nutrient-dependent regulation of translation rather than regulated histone stability that compensates for the transcriptional downregulation in poor nutrients.

Given the importance of the 5’ untranslated region (UTR) in controlling translation efficiency and protein abundance we sought to examine the influence of the *HTB1* 5’ UTR on the nutrient-dependent mCitrine expression from the *HTB1* promoter. To this end, we replaced the *HTB1* 5’ UTR with the 5’ UTRs of *MDN1* and *RPB4*, respectively (Fig. 5D, Supplementary Fig. 7C, D Supplementary Table 4). *RPB4* is a subunit of the RNA polymerase II complex and its expression has been reported to scale with cell size (*45*). If sequences within the *HTB1* 5’ UTR contribute to maintaining the histone protein amounts constant in rich and poor medium, then these replacements could lead to a change in the protein regulation. First, we performed RT-qPCR and found that replacing the 5’ UTR had no significant effect on the nutrient-dependence of the *mCitrine* mRNA concentrations, which were decreased in nutrient-poor conditions (Fig. 5E). Moreover, while flow cytometry analysis of mCitrine protein expression revealed an overall decrease in the protein abundance after substitution of the *HTB1* 5’-UTR, the protein amounts remained constant between rich and poor conditions (Fig. 5F). This suggests that, despite its influence on protein abundance, the histone 5’ UTR is not required for nutrient-dependent histone homeostasis.

### Cells are more sensitive to excess histone accumulation under non-fermentable growth conditions

So far, we have characterized nutrient-dependent histone homeostasis at protein and transcript levels. We showed that cells maintain constant stoichiometry between DNA and histone proteins. Surprisingly, histone transcripts are less abundant in poor nutrient conditions, raising the question of why it could be beneficial for cells to downregulate histone transcripts even though protein amounts are maintained constant. Interestingly, a recent study showed that cells exhibit a lower tolerance to excess histones under glucose-limited conditions, resulting in decreased cell fitness (*46*). In our case, this could imply that cells in poor, *i*.*e*. non-fermentable, growth medium keep the histone transcript concentrations low to reduce the risk of accumulating excess histones. To assess the importance of nutrient-dependent transcript regulation, we tested how cells respond to aberrantly high levels of histone transcripts under fermentable and non-fermentable growth conditions. For this purpose, we followed the strategy of Bruhn et al. (*46*), and used a *rad53Δ* mutant, which is defective in histone transcript regulation and in the degradation of excess histones (Fig. 6A). Cells lacking Rad53, a DNA damage response kinase, are only rendered viable by additional deletion of the ribonucleotide reductase inhibitor Sml1. The *sml1Δ* mutant has a similar doubling time and cell volume compared to the wild type and served as a reference (Fig. 6B, Supplementary Fig. 7E, F). Analysis of the histone transcript concentrations in the *sml1Δrad53Δ* mutant revealed increased accumulation of *HTB2* and *HHT2* transcripts compared to the *sml1Δ* strain, on both rich and poor carbon source (Fig. 6C, D). We included *HHT2* in our analysis as a control, as it has been already shown by Bruhn et al. (*46*) to exhibit elevated mRNA expression concentrations in *sml1Δrad53Δ* cells. However, *HTB1* transcript concentrations were not affected by deletion of *RAD53*. Despite that, western blot analysis revealed that *sml1Δrad53Δ* mutants have increased concentrations of H2B protein in both growth media (Fig. 6E). To examine whether this overexpression affects cell growth on rich and poor carbon source, we measured the corresponding doubling times (Fig. 6B). In SCGE, we observe that *sml1Δrad53Δ* cells grow significantly slower than the reference *sml1Δ* strain. In YPD, however, the lack of Rad53 does not cause major changes in the doubling time. Given that *sml1Δrad53Δ* mutants are also defective in DNA replication (*47*–*49*) and DNA damage response (*48, 50, 51*), among others, we sought to disentangle the contribution of excess histone accumulation to the observed growth phenotype. For this purpose, we drastically decreased histone protein levels by deleting *SPT21*, a histone transcription activator, and examined the effects on cell growth. We find that in SCGE, *sml1Δrad53Δspt21Δ* triple deletion mutants grow faster, indicating that the growth phenotype of *sml1Δrad53Δ* cells is partly rescued. In contrast, the doubling time in YPD is not noticeably affected (Fig. 6B). Thus, our results suggest that not only upon glucose limitation (*46*) but also during growth on non-fermentable carbon sources, cells exhibit higher sensitivity to excess histone accumulation than in rich glucose conditions. This suggests that on poor nutrients, histone transcription is downregulated to avoid toxic overexpression, but in rich nutrients rapid histone production becomes more important. To further test this hypothesis, we created a diploid strain in which we deleted the endogenous alleles of *HTB2* and one allele of *HTB1* to examine the effects of decreased histone amounts on cell growth in rich and poor conditions. While cells in SCGE are unaffected in their doubling time, in YPD we find that reducing the histone amounts leads to slower cell growth (Supplementary Fig. 8).

Taken together, our results suggest that nutrient-dependent regulation of histone transcripts is important because – depending on the nutrient condition – histone overexpression affects cells to different degrees, which shifts the optimal balance between production speed and prevention of excess production.

## Discussion

Despite changes in cell growth and cell cycle progression, cells maintain constant histone-to-DNA stoichiometry across different nutrient conditions. This implies that histone protein concentrations are higher in poor compared to rich nutrients, to account for the smaller cell size. Paradoxically, histone transcripts show the opposite trend, and are downregulated in poor nutrients. The apparent decoupling of histone mRNA and protein nutrient-dependence suggests that histone-specific nutrient-dependent regulation of translation is required to ensure constant histone amounts.

Previous work on cell-volume dependent histone homeostasis showed that the coordination of histones with genome content is already established at the transcript level and is at least in part mediated by the promoter (*31, 32*). This differential regulation of histones was proposed to be achieved through template-limited transcription, where the gene itself, rather than the polymerase (*45*), limits histone mRNA synthesis (*31*). Our single cell analysis revealed that for each nutrient condition, transcript amounts expressed from a histone promoter are indeed independent from cell volume. However, the fact that transcript amounts decrease significantly in poor compared to rich growth media, suggests that nutrient-dependent histone regulation cannot be explained by ‘gene-limited transcription’ alone. Highlighting this distinct regulation of histones, we found that in contrast to histone mRNAs, the mRNA amounts of *ACT1* and *MDN1*, two representatives of scaling gene expression, increase in proportion to cell volume, but are largely independent of the nutrient condition.

As histone synthesis is restricted to S-phase, it could in principle be possible that nutrient-induced changes in the relative duration of the cell cycle phases explain the decrease of histone transcripts in poor growth media. Cells growing on SCGE spend more time in G1 compared to growth on YPD, resulting in relatively shorter S- and G2/M-phases. Specifically, cell cycle analysis by flow cytometry revealed that the fraction of S-phase cells decreases from 36.4 % in YPD to 29.8% in SCGE (Fig.1D). This relatively moderate difference of 7% suggests that the nutrient-dependent change of the relative S-phase duration alone cannot explain the downregulation of histone transcripts in SCGE by a factor of two compared to YPD. Ultimately, our findings suggest that nutrient-dependent transcription must account for the reduced transcript levels, as mRNA stability increases in poor conditions.

Why do cells down-regulate histone transcripts if the amount of protein needs to be coordinated with the DNA content and therefore is kept constant across nutrient conditions? Previously, it has been shown that upon glucose limitation, cells are more sensitive to histone overexpression. This sensitivity is partly induced by the hyper-acetylation of excess histones, which under poor growth conditions affects the Ac-CoA-dependent metabolism to a greater extent, due to the lower availability of Ac-CoA (*46*). We now show that also for cells growing on non-fermentable carbon sources histone overexpression is more toxic than for cells grown on glucose. Our findings therefore suggest that cells growing on poor nutrients reduce the risk of accumulating excess histones by maintaining low concentrations of histone transcripts. In rich nutrient environments, on the other hand, where cells exhibit higher growth rates, fast histone production may be more critical.

The protein-to-mRNA ratio is dictated by translation and protein degradation. Thus, the uncoupling of the regulation of protein and mRNA abundances we observed in different nutrients implies the need for nutrient-specific regulation of translation or protein stability. Specifically, our data suggest that to compensate for the decreased transcript concentrations in poor nutrients, the relative translation efficiency, or stability, of core histones is higher than in rich nutrients.

We have shown that histone promoters mediate nutrient-dependent transcription as well as its compensation on the protein level even when expressing the fluorescent reporter mCitrine. This indicates that the decoupling of histone transcript and protein nutrient-regulation is achieved mainly through nutrient-dependent translation rather than protein degradation. Interestingly, our experiments revealed that the histone 5’ UTR is not required for maintaining constant amounts of proteins expressed from a histone promoter across changing nutrients. This indicates that histone translation is regulated through an “imprinting” mechanism: Previous studies have identified several factors that bind to newly produced mRNAs in the nucleus and remain associated with them throughout their lifecycle, controlling mRNA export, localization, decay or translation (*52*). For example, it was proposed that Pol II remotely modulates mRNA translation and decay through co-transcriptional binding of the Rpb4/7 heterodimer to Pol II transcripts (*53*–*55*). Thereby in addition to the encoded information, mRNAs would carry information “imprinted” by Rpb4/7, which is required for proper post-transcriptional regulation. Moreover, several studies of promoter-dependent mRNA stability suggest that promoter elements can also contribute to mRNA imprinting providing cross talk with cytoplasmic processes (*37, 56*).

Overall, we have characterized the distinct regulation of histone homeostasis in changing environments, highlighting the importance for cell-cycle dependent genes to maintain accurate protein concentrations despite the nutrient-induced changes in cell growth and cell cycle progression. Our work revealed a surprising mode of regulation, where histone protein concentrations are decoupled from transcript concentrations. We speculate that this allows cells to balance the need for rapid production under fast growth conditions with the tight regulation required to avoid toxic overexpression in poor media (Fig. 7). More generally, this suggests that cells use separate regulation of transcripts and translation as a way to not only control the final protein concentration but also optimize the balance between production speed and accuracy. Future studies will be needed to reveal whether such regulation also occurs for other genes, in particular cell cycle dependent genes that are high expressed only during a short fraction of the cell cycle.

## Materials and Methods

### Strains and culture conditions

Budding yeast strains used in this study are haploid derivatives of W303 and were constructed using standard procedures. All transformants were validated using PCR and sequencing. Full genotypes for each strain can be found in Supplementary Table 1.

Cells were grown under different nutrient conditions at 30 °C in a shaking incubator at 250 rpm (Infors, Ecotron). Yeast colonies were inoculated in 4 mL yeast peptone medium containing 2% glucose (YPD) and were cultivated for at least 6 h at 30 °C before being washed and transferred to synthetic complete (SC), yeast peptone (YP) or minimal medium (SD), with either glucose (D) or galactose (Gal) as fermentable or glycerol and ethanol (GE) as non-fermentable carbon sources. Cells were then grown in the respective growth medium for at least 18 h to OD_600_ = 0.3 – 0.9. Through appropriate dilutions, cell density was maintained below OD_600_ = 1. Optical densities were measured using a spectrophotometer (NanoDrop One^C^, Thermo Fisher Scientific).

### Western blot

Total protein extracts were prepared according to a previously established protocol (*57*). Briefly, cell cultures (25 mL) were grown in different growth media for at least 18 h to ensure steady-state conditions (see above). Prior to harvesting, cell volume distributions and cell numbers per mL were determined using a Coulter Counter (Beckman Coulter, Z2 Particle Counter). From each culture, 5×10^7^ cells were collected by centrifugation (4k rpm, 3 minutes) and washed with 1 mL of ice-cold double-distilled water before being spun down (10k rpm, 2 minutes) and resuspended in 400 μL of 0.1 M NaOH. After incubation at room temperature (RT) for 10 min, cells were again pelleted (10k rpm, 2 minutes), and then boiled (3 min, at 95 °C) in 120 μL of reducing 1x LDS sample buffer, containing 30 μL of 4X Bolt™ LDS sample buffer (Invitrogen), 12 μL of 10X Bolt™ sample reducing agent (Invitrogen) and 78 μL of double-distilled water.

Following centrifugation (10k rpm, 2 minutes), 5-10 μL of total protein extracts (supernatant) were loaded into each lane of commercially available Bolt™ 12% Bis-Tris plus mini-gels (Invitrogen). Gels were run (200V, 160 mA, 20-25 min) in 1X Bolt™ MES SDS Running Buffer (Invitrogen) and the separated proteins were then transferred onto nitrocellulose membranes (10 V, 160 mA, 60 min) using the mini-blot-module (Invitrogen). In a next step, membranes were stained with Ponceau S and total proteins were visualized using the ChemiDoc™ MP imaging system (Bio-Rad). To detect proteins of interest, membranes were blocked in TBST (Tris-buffered saline, 0.2% Tween 20) with 5% milk and incubated overnight at 4 °C with the primary antibodies: rabbit monoclonal anti-histone H2B (Abcam Cat#ab188291, 1:2000) or mouse monoclonal anti-beta actin (Abcam, Cat# ab170325, 1:10000). After washing the membranes in TBST (3 × 5 min), they were probed (1.5 h, RT) with the HRP-conjugated secondary antibodies: goat anti-mouse IgG (Abcam Cat# ab205719, 1:10000) or goat anti-rabbit IgG (Abcam Cat# ab205718, 1:10000). To visualize the protein bands, membranes were incubated (5 min, RT) in Clarity™ western ECL substrate (Bio-Rad) and imaged using the ChemiDoc™ MP imaging system (Bio-Rad). Quantification of band intensities was carried out using the Image Lab 5.2.1 software (Bio-Rad).

### RNA extraction and RT-qPCR

Total RNA was extracted from 2-5 x10^7^ cells grown in different nutrients (see above) with the YeaStar RNA Kit (Zymo Research) following the manufacturer’s instructions. Prior to harvesting, cell numbers per mL were measured using a Coulter Counter. Concentration and purity of the eluted RNA were determined with a spectrophotometer (NanoDrop One^C^, Thermo Fisher Scientific) before 800 ng total RNA was reverse transcribed using random primers and the high-capacity cDNA reverse transcription kit (Thermo Fisher Scientific). To measure relative mRNA levels of target genes, the obtained cDNA was diluted 10-fold (*HTB2, ACT1, MDN1, mCitrine*) or 100-fold (*HTB1, HTA1, HTA2, HHT1, HHT2, HHF1, HHF2*) in double-distilled water and 2 μL of the dilutions were used as templates for quantitative PCR (qPCR). All qPCR reactions were performed on a LightCycler 480 Multiwell Plate 96 (Roche) using the SsoAdvanced Universal SYBR Green Supermix (BioRad,) and target-specific primers (Supplementary Table 2). For each target gene, mean Cq^Gene^ values of three technical replicates per sample were normalized to the reference gene *RDN18*, and relative mRNA concentrations were calculated by the formula: log_2_ (relative concentration) = − (Cq^*Gene*^ – Cq^*RDN18*^).

### mRNA stability measurements

Cell cultures (50 mL) were grown in YPD and SCGE, respectively, for at least 18 h to OD_600_ = 0.3 – 0.5 (see above). To determine the half-lives of the *HTB1, HTB2* and *ACT1* mRNAs, cells were treated with the RNA polymerase inhibitor thiolutin (Biomol) at a final concentration of 8 μg/mL(*58*). After the addition of thiolutin, 4 mL samples were taken at given time points from 0 to 60 min of incubation. Cells were harvested by centrifugation (2500 x g, 3.5 min) and washed with 1 mL of RNAse-free water (Qiagen) before being spun down (10k rpm, 2 minutes) and resuspended in 80 μL of digestion buffer (included in the YeaStar RNA Kit, Zymo Research). Cells were then stored on ice until ready for RNA extraction, which was performed using the YeaStar RNA Kit (Zymo Research). To remove DNA contaminations, the RNA samples were treated with DNAse I (Life Technologies). Relative changes in mRNA concentrations were measured by RT-qPCR as described above. Target-specific primers were designed to bind to the *HTB1, HTB2* and *ACT1* coding sequences, respectively (*36*). The half-lives of histone and *ACT1* mRNAs were calculated by fitting a single exponential function to the obtained mRNA decay curves.

### Flow Cytometry

Cells (2-5 mL) were cultured at 30 °C in different growth media for 36 h. During growth, cell cultures were kept at OD_600_ < 1. For each sample, cell volume distributions were measured using a Coulter Counter and bud counts were performed to estimate the fraction of budded and unbudded cells in the populations. To analyse the fluorescence intensity of mCitrine expressed in wildtype cells, flow cytometry measurements were carried out on a 577 CytoFlex S Flow Cytometer (Beckman Coulter). mCitrine was excited with a 488-nm laser and detected using a 525/40-nm bandpass filter. 50,000 events were analysed in each experiment at a flow rate of 10μL/min, which corresponds to roughly 1000 events/sec. Manual gating based on the side scatter (SSC) and forward scatter (FSC) parameters was performed using the FlowJo 10.8.1 software (Becton Dickinson, San Josè, CA) to eliminate doublets and cell debris (Supplementary Fig. 2A). Wildtype cells not expressing mCitrine were analysed in all conditions to correct for autofluorescence.

Flow cytometry was also applied to determine the cell cycle distribution of wildtype cells in different nutrients by quantification of the DNA content. For this purpose, cells were fixed and stained with the fluorescent DNA-binding dye SYBR Green I, according to a previously published protocol (*59*). Briefly, 1 mL of cell culture grown for 36 h in the respective growth medium (final OD_600_ = 0.5) was slowly added to 9 mL of 80 % ethanol and incubated overnight at 4 °C. Next, cells were pelleted (2.500 x g, 2 min, 4°C) and washed twice with 50 mM Tris-HCl (pH = 8.0) before being treated with 300 μL of 1 mg/mL RNase A at 37°C for 40 min. After washing with 50 mM Tris-HCl (pH = 8.0), cells were incubated in 50 μL 20 mg/mL Proteinase K at 37°C for 60 min. In a last step, cells were washed again with 50 mM Tris-HCl (pH = 8.0) and treated with 200 μL of 10x SYBR Green I (Sigma-Aldrich) DNA stain at 22°C for 1 h. For quantification of the cellular DNA content, SYBR Green I was excited with a 488-nm laser and detected using a 525/40-nm bandpass filter. The obtained DNA frequency histograms showed defined G1 and G2 peaks and were analysed with the FlowJo 10.8.1 software (Becton Dickinson, San Josè, CA). The Watson pragmatic algorithm was used to model the cell cycle and estimate the percentages of cells in the different cell cycle phases.

SYBR Green I is not suitable for staining the DNA of cells expressing mCitrine due to the overlapping emission spectra of the two fluorophores. However, cells with *mCitrine*-tagged H2B also show fluorescence histograms with distinct G1 and G2 peaks, since core histone synthesis is tightly coupled to the DNA replication during S-phase. Thus, live cells were measured as described above and the obtained fluorescence histograms of mCitrine were analyzed in order to determine the cell cycle distributions in different nutrient conditions.

### Single-molecule fluorescence in situ hybridization (smFISH)

For smFISH analysis, commercially available Stellaris® FISH probes were designed using the Stellaris® FISH Probe Designer (Biosearch Technologies). More precisely, the, *MDN1* and *mCitrine* transcripts were targeted with probe sets consisting of 27 to 48 20-mer oligonucleotides labeled with the dye Quasar-670^®^, while *ACT1* transcripts were bound by 41 individual Quasar-570^®^-labeled probes.

smFISH samples were prepared following the Stellaris® RNA FISH protocol for *S. cerevisiae*, available online at www.biosearchtech.com/stellarisprotocols. 45 ml cultures were grown in different growth media for at least 18 h to an OD_600_ = 0.3-0.5 before being fixed for 45 min at room temperature by adding formaldehyde to a final concentration of 4%. After centrifugation (1600 x g, 4 min), cells were washed twice with 1 mL of ice-cold fixation buffer (1.2 M sorbitol (Sigma-Aldrich), 0.1 M K2HPO4 (Sigma-Aldrich), pH 7.5) and incubated at 30 °C for 55 min in 1 mL of fixation buffer containing 6.25 μg zymolyase (Biomol). Digested cells were then washed twice with 1 mL of ice-cold fixation buffer and stored overnight at 4 °C in 1 mL of 70% ethanol. Next, 300 uL of cells were spun down and hybridized overnight at 30 °C with 100 uL of Stellaris® RNA FISH hybridization buffer (Biosearch Technologies) containing 10% v/v formamide and 125 mM smFISH probes. The following morning, cells were washed with 10% v/v formamide in Stellaris® RNA FISH wash buffer A (Biosearch Technologies) and incubated in 1 mL of DAPI staining solution (5 ng/mL DAPI in wash buffer A with 10% v/v formamide) for 30 min at 30 °C. After washing with 1 mL of Stellaris® RNA FISH wash buffer B (Biosearch Technologies) cells were mounted in Vectashield® mounting medium (Vector Laboratories). Wide-field fluorescence imaging was performed using a Zeiss LSM 800 microscope equipped with a 63×/1.4 NA oil immersion objective and an Axiocam 506 camera. Multicolor z-stacks composed of 20 images were recorded at 240 nm intervals with the Zen 2.3 software. Quasar-570^®^ and Quasar-670^®^ were illuminated with a 530 nm LED and a 630 nm LED respectively, while DAPI images were taken using a 385 nm LED.

### smFISH image analysis

Cell segmentation was performed using Cell-ACDC (*60*). Briefly, cells were segmented based on bright-field signal using YeaZ (*61*) and buds were manually assigned to the correct mother cells. Cell volumes were automatically calculated by Cell-ACDC starting from the generated 2D segmentation masks. To detect and count the number of fluorescence spots in 3D we developed a custom routine written in Python. The analysis steps are the following: 1) Application of a 3D gaussian filter with a small sigma (0.75 voxel) of both the nucleoids and mitochondria signals. 2) Instance segmentation of the spots’ signal using the best-suited automatic thresholding algorithm (either the threshold triangle, Li, or Otsu algorithms from the Python library scikit-image (*62*)). 3) 3D local maxima detection (peaks) in the spots signal using the *peak_local_max* function from the Python library scikit-image. Note that peaks are searched only on the segmentation masks determined in the previous step. 4) Discarding of overlapping peaks: if two or more peaks are within a resolution-limited volume, only the peak with highest intensity is retained. The resolution-limited volume is determined as a spheroid with x and y radii equal to the Abbe diffraction limit and z radius equal to 1 μm. For example, with a numerical aperture of 1.4 and Quasar-670^®^ emission wavelength of about 668 nm, the resolution limited volume has x=y=0.291 μm radius. 5) The remaining peaks undergo a subsequent iterative filtering routine: for each peak, we computed the Glass’ delta (a measure of the effect size) of the peak’s pixels compared to the background’s pixels. The pixels belonging to the peaks are defined as the pixels inside the resolution-limited volume explained in step 4. The pixels belonging to the background are those pixels outside of all the detected peaks but inside the segmentation mask of the cell. The Glass’s delta is computed as the mean of the peak’s signal (positive signal) subtracted by the mean of the background’s signal (negative or control signal) all divided by the standard deviation of the background’s signal. Peaks that have an effect size lower than a threshold are discarded. The threshold value was manually determined for each experiment after careful inspection of the images. This value ranged from 0.2 to 1.0. Finally, step 5 was repeated until the number of peaks stopped changing. The final number of peaks corresponds to the detected mRNA spots.

To determine the cell cycle stage, the cells were grouped into either G1-, S- or G2/M-phase based on the bud-to-mother cell volume ratio, as well as the number of DAPI-stained nuclei in the cell. Unbudded cells containing one nucleus were classified as G1 cells. If a bud was present, the ratio of bud volume divided by mother volume was used as a classification criterion. Cells with one nucleus and a ratio < 0.3 were considered to be in S-phase. For ratios > 0.3, cells were classified as G2/M cells (*31*). Lastly, if cells were budded and contained two nuclei, cell outlines were carefully inspected in the bright-field images to distinguish G2/M cells from two neighboring G1 cells. mRNA concentrations were estimated as the number of mRNA spots per cell divided by the cell volume

### Live-cell fluorescence microscopy

Cells, pre-cultured in YPD, were transferred to a selected growth medium and were grown for at least 18 h to OD_600_ = 0.3 – 0.9. For live-cell imaging, 200 μL of cells were sonicated for 5 s before being loaded into a CellASIC® ONIX2 Y04C microfluidic plate (Y04C, Millipore) connected to the ONIX2 (Millipore) microfluidic pump system. Cells were trapped inside the microfluidic chamber and fresh medium was continuously supplied with a constant pressure of 13.8 kPa. Live-cell fluorescence microscopy was performed on a Zeiss LSM 800 microscope equipped with an epifluorescence setup and coupled to an Axiocam 506 camera. Cells were imaged in a 30 °C incubation chamber (Incubator XLmulti S1, Pecon) at a 3 min interval over the course of 7 to 12 hours using an automated stage (WSB Piezo Drive Can) and a 40×/1.3 NA oil immersion objective. Cells with *mCitrine*-tagged *HTB1* and/or *HTB2* were illuminated for 10 ms with a 511 nm LED set at 5% power. To measure mCitrine expression from the *HTB1, HTB2* or *ACT1* promoter, the LED power was increased to 12% due to the lower emission intensity of mCitrine in those cells.

For analysis, the collected microscopy images were aligned and cropped to the region of interest using a custom Fiji script (*63*). Cell segmentation and tracking were performed as previously described by Doncic et al (*64*). Briefly, cells were segmented based on the phase contrast images and tracked backwards in time. Pedigrees were then manually annotated, and the time points of cell birth, bud emergence and division were identified through visual inspection for all daughter cells born during the time-lapse experiment. The total fluorescence measured per cell was background-corrected as described by Chandler-Brown et al (*65*). Moreover, for quantification of the fluorescence intensity of mCitrine expressed from the *HTB1, HTB2*, or *ACT1* promoter, nutrient-and cell-volume-dependent autofluorescence was subtracted (*65*). Autofluorescence was determined by analyzing wildtype cells not expressing a fluorescent protein.

In the case of *mCitrine*-labeled histones, autofluorescence is much weaker than the mCitrine fluorescence signal in all nutrient conditions (Supplementary Fig. 3D), and was therefore neglected. For the control experiments shown in Supplementary Fig. 3D, cell segmentation and quantification of the total fluorescence signal per cell were perfomed using Cell-ACDC (*60*).

For all cells born during the experiment, fluorescence intensity traces were analyzed to quantify mCitrine dynamics during the cell cycle. The single cell expression profile of the endogenously tagged histones shows a plateau during G1, followed by a 2-fold increase in fluorescence starting around bud emergence and a second plateau during G2/M-phase. The total amount of mCitrine produced during the cell cycle was calculated as the difference between the median of the first four time points and the median of the last four time points.

### Statistical Analysis

Statistical analysis was performed using GraphPad Prism 9.4.1 All data sets were tested for normality using the Shapiro–Wilk test at a confidence level of α = 0.05. Significances were calculated using an unpaired, two-tailed t-test for datasets that follow a Gaussian distribution or a Mann-Whitney test for datasets that are not normally distributed. To compare the slopes of two linear regression lines, p-values were computed using GraphPad Prism 9.4.1.

## Acknowledgements

We thank Christopher Bruhn and Marco Foiani for sharing strains, Pascal Falter-Braun, Christof Osman and members of the Institute of Functional Epigenetics for insightful scientific discussions. We thank Matthew Swaffer for helpful comments on the manuscript. This work was funded by the Deutsche Forschungsgemeinschaft (DFG, German Research Foundation) through projects 416098229 and 431480687, by the Human Frontier Science Program (career development award to K.M.S.), and the Helmholtz Association.

