## Supplementary Information for "Decoupling of transcript and protein concentrations ensures budding yeast histone homeostasis in different nutrient conditions"

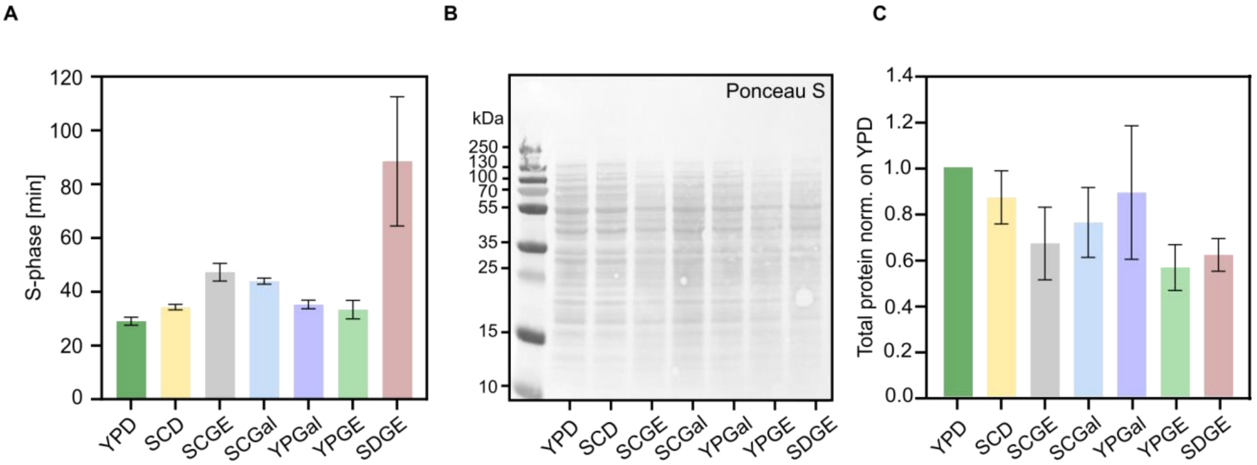


**Supplementary figure 1**. **Nutrient-dependent changes in S-phase duration and total protein abundance.** (**A**) Absolute S-phase durations in different nutrient conditions were calculated from the cell cycle fractions shown in Fig. 1D using the corresponding population doubling times. Bar plots with error bars indicate mean values and standard errors across n = 5 replicate measurements. (**B**) Representative western blot membrane stained with Ponceau S for quantification of total proteins. (**C**) Total protein content, extracted from equal number of cells in different growth media, was quantified by Ponceau S staining and normalized on YPD. Bar plots with error bars indicate mean values and standard deviations of at least 4 independent replicates.

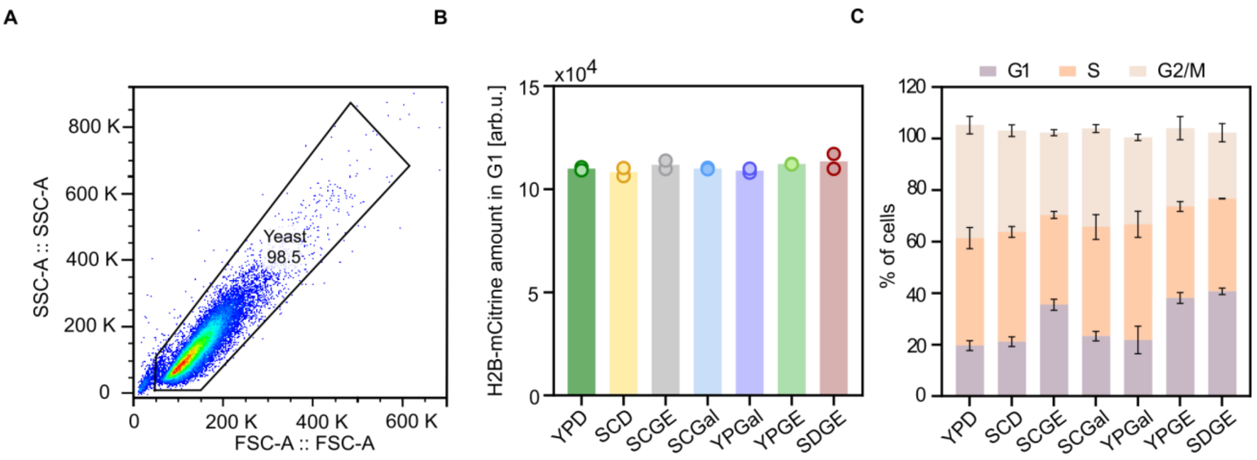


**Supplementary figure 2**. **Flow cytometry experiments reveal constant histone amounts during G1 in all nutrient conditions.** (**A**) Example of gating strategy used in flow cytometry analysis. (**B**) Total H2B-mCitrine (Htb1 and Htb2 tagged) amounts in G1 were quantified using flow cytometry in different growth media. The bar graphs represent the mean of n = 2 independent replicates, each shown as an individual dot. (**C**) Nutrient-dependent cell cycle distributions (percentage of cells in G1-, S- and G2/M-phase) of cells with *mCitrine*-tagged H2B, determined with flow cytometry using the H2B fluorescence intensity. Error bars represent the standard deviation of n = 2 independent measurements.

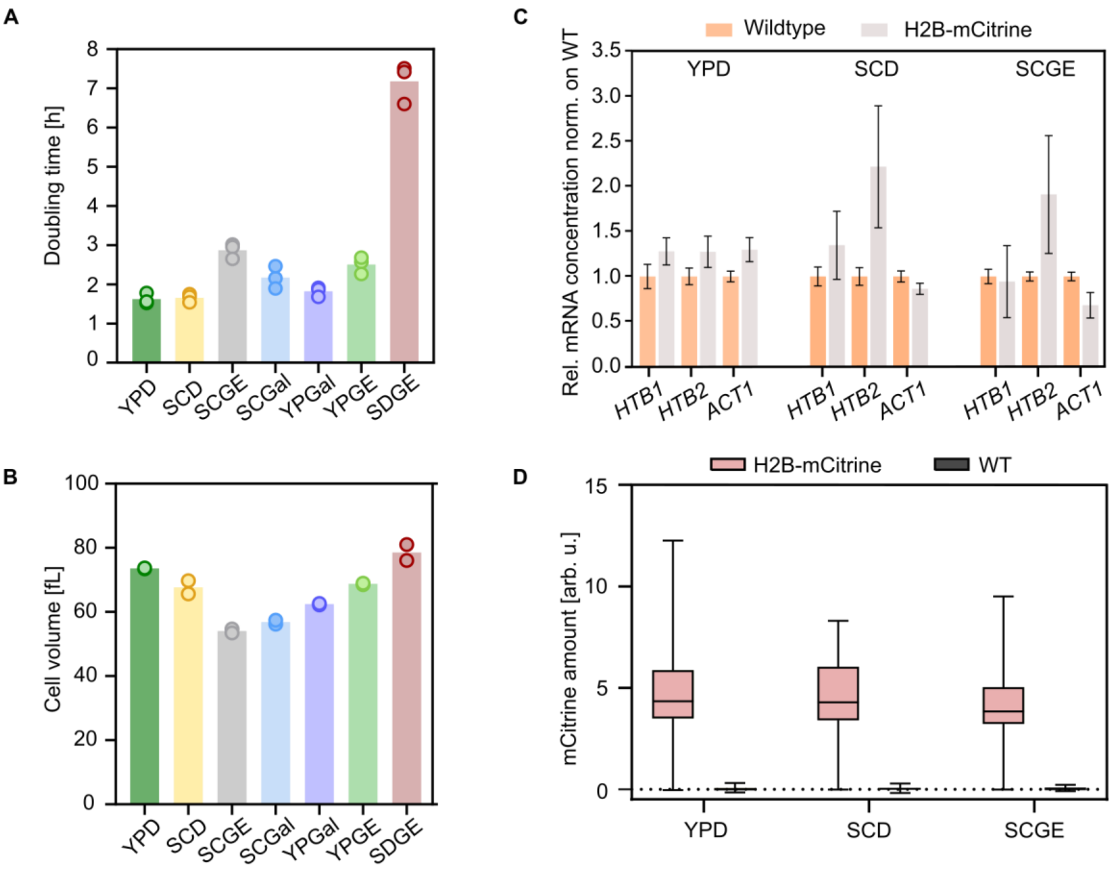


**Supplementary figure 3. Characterization of H2B-mCitrine strain in different growth media.** (**A**) Doubling times calculated from growth curves of cells with *mCitrine*-tagged *HTB1* and *HTB2,* growing in different nutrient conditions. Bars represent the mean of n = 3 independent measurements shown as individual dots. (**B**) Corresponding nutrient-specific mean cell volumes measured with a Coulter counter. Bars represent the mean of n = 2 independent measurements shown as individual dots. (**C**) RT-qPCR was used to measure the mRNA concentrations of *HTB1* and *HTB2*, as well as the control gene *ACT1* in wildtype cells and cells with *mCitrine*-tagged *HTB1* and *HTB2*. mRNA concentrations were normalized on *RDN18* and are shown as fold changes compared to the wildtype strain. Bars represent the mean values of at least 3 independent biological replicates; error bars indicate standard errors. (**D**) Compared to H2B-mCitrine intensity, autofluorescence is negligible. Total mCitrine fluorescence intensity after background correction was measured in cells with mCitrine-tagged H2B ($n_{YPD}$ = 492, $n_{SCD}$ = 392 $n_{SCGE}$ = 275) and untagged H2B ($n_{YPD}$ = 227, $n_{SCD}$ = 285 $n_{SCGE}$ = 215). Box plots represent median and 25th and 75th percentiles, whiskers are extending to the minimum and maximum values.

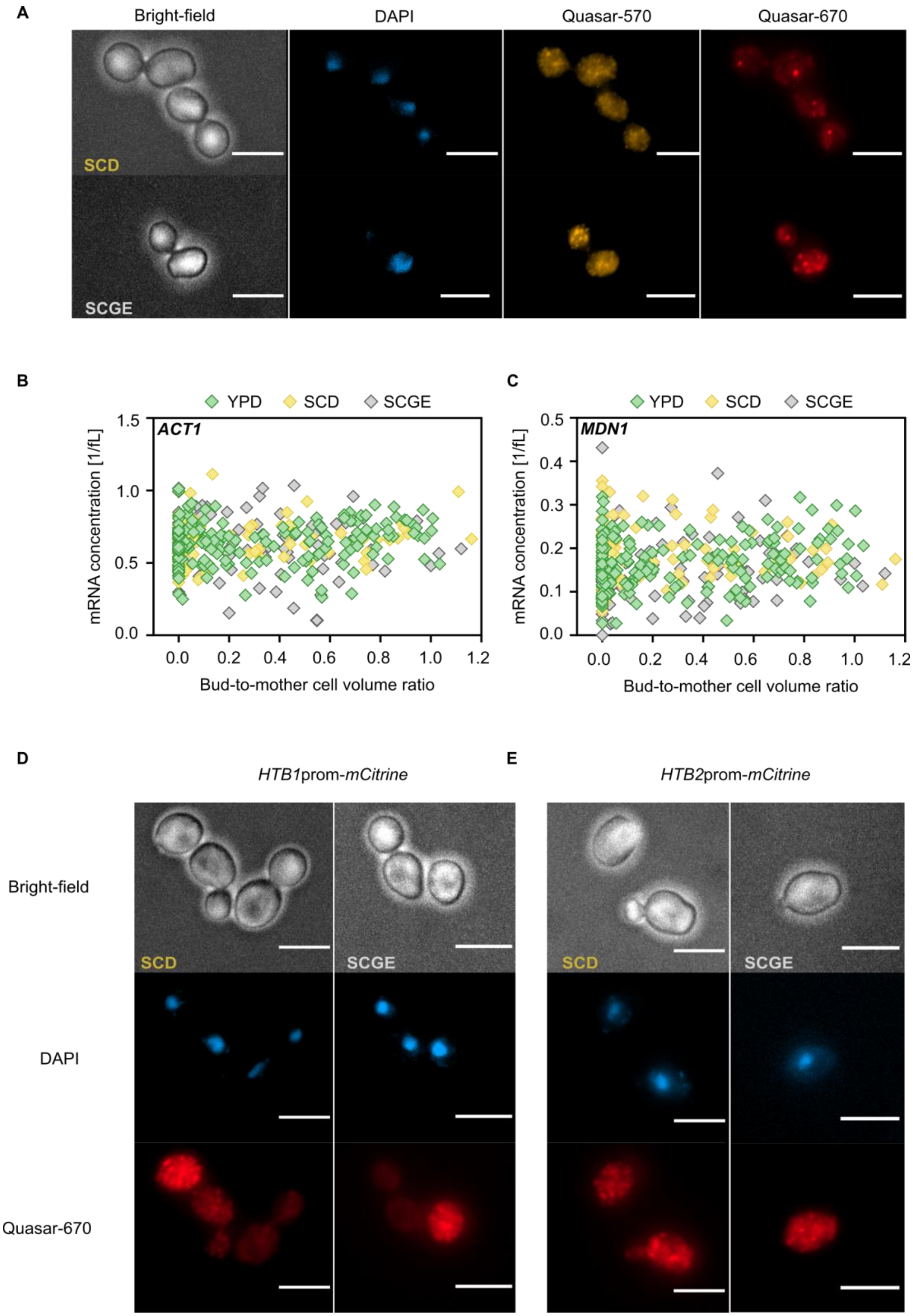
A
Bright-field
DAPI
Quasar-570
Quasar-670
SCD
SCGE
B
YPD
SCD
SCGE
ACT1
mRNA concentration [1/fL]
1.5
1.0
0.5
0.0
0.0
0.2
0.4
0.6
0.8
1.0
1.2
Bud-to-mother cell volume ratio
C
YPD
SCD
SCGE
MDN1
mRNA concentration [1/fL]
0.5
0.4
0.3
0.2
0.1
0.0
0.0
0.2
0.4
0.6
0.8
1.0
1.2
Bud-to-mother cell volume ratio
D
HTB1prom-mCitrine
E
HTB2prom-mCitrine
Bright-field
DAPI
Quasar-670
SCD
SCGE
SCD
SCGE

**Supplementary figure 4. *ACT1* and *MDN1* are expressed throughout the cell cycle at constant, nutrient-independent concentrations.** (**A**) *ACT1* and *MDN1* transcripts were detected in single cells using smFISH probes labeled with Quasar-570 (yellow) and Quasar-670 (red), respectively. Nuclear DNA was stained with DAPI (blue). Representative images of cells grown in SCD and SCGE are shown. The scale bars represent 5 μm. (**B-C**) mRNA concentrations (mRNA spots per cell volume) of (**B**) *ACT1* ($n_{YPD}$ = 176, $n_{SCD}$ = 87, $n_{SCGE}$ = 98) and (**C**) *MDN1* ($n_{YPD}$ = 176, $n_{SCD}$ = 87, $n_{SCGE}$ = 98) were plotted against the corresponding bud-to-mother cell volume ratio in different nutrients. (**D-E**) Representative images of cells expressing (**D**) *HTB1prom-mCitrine* and (**E**) *HTB2prom-mCitrine* in SCD and SCGE. *mCitrine* transcripts were detected using probes labeled with Quasar-670 (red). Cell nuclei were stained with DAPI (blue). The scale bars represent 5 μm.

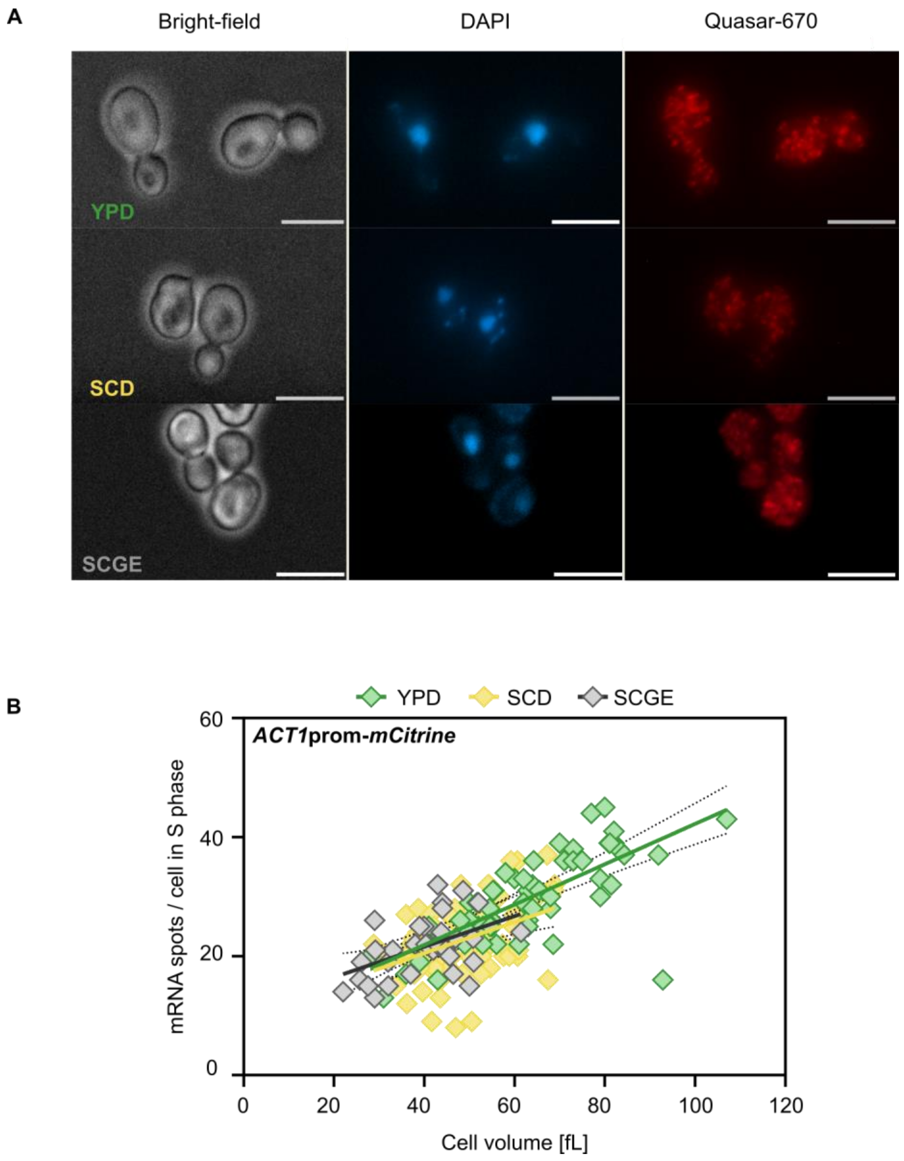


**Supplementary figure 5. Transcript amounts expressed from *ACT1* promoter increase with cell volume in all nutrient conditions.** (**A**) Representative images of cells expressing *ACT1*prom-*mCitrine* in YPD, SCD and SCGE. *mCitrine* transcripts were detected using probes labeled with Quasar-670 (red). Cell nuclei were stained with DAPI (blue). The scale bars represent 5 µm. (**B**) Number of *mCitrine* mRNA spots per cell (*ACT1*promoter-*mCitrine*; $n_{YPD}$ = 59, $n_{SCD}$ = 75, $n_{SCGE}$ = 33) as a function of cell volume during S-phase. Here, cells with one nucleus and a bud-to-mother volume ratio < 0.3 were considered to be in S-phase. Lines show linear fits; dashed lines indicate the 95% confidence intervals.

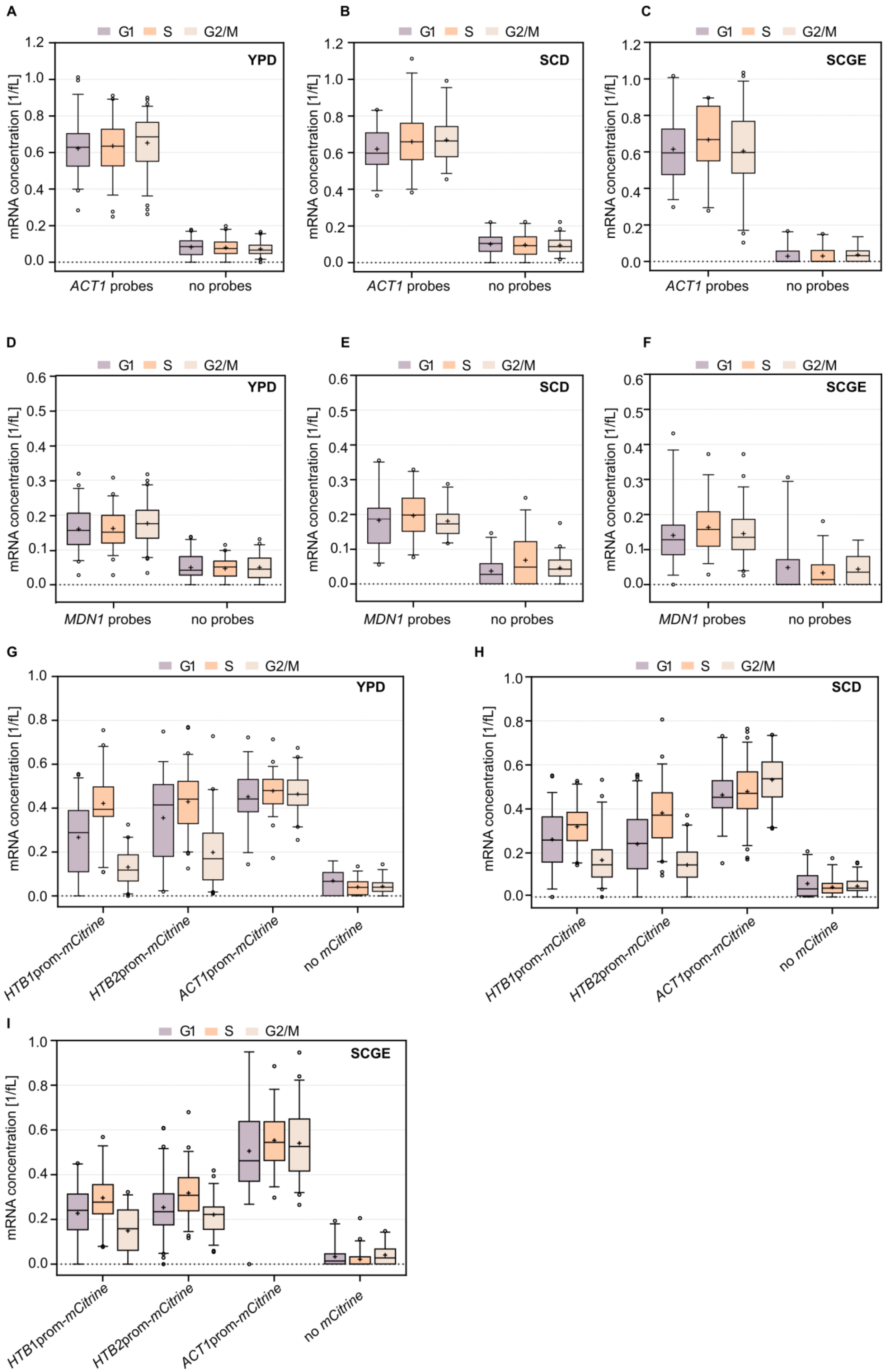
A
B
C
D
E
F
G
H
I
G1
S
G2/M
YPD
SCD
SCGE
mRNA concentration [1/fL]
ACT1 probes
MDN1 probes
no probes
HTB1prom-mCitrine
HTB2prom-mCitrine
ACT1prom-mCitrine
no mCitrine

**Supplementary figure 6. mRNA concentration of transcripts of interest in G1-, S- and G2/M-phase measured with smFISH are shown in comparison to negative controls.** Wildtype cells were incubated with or without smFISH probes against (**A-C**) *ACT1,* (**D-F**) and *MDN1*. mRNA concentrations in G1, S and G2/M were estimated by dividing the detected spots by the cell volume. (**G-I**) Wildtype cells expressing no mCitrine*,* as well as cells carrying an additional copy of the *HTB1, HTB2* or *ACT1* promoter driving mCitrine were incubated with smFISH probes against *mCitrine*. mRNA concentrations in G1, S and G2/M were estimated by dividing the detected spots by the cell volume. Box plots represent median and 25th and 75th percentiles; whiskers indicate the 5th and 95th percentiles and symbols show outliers.

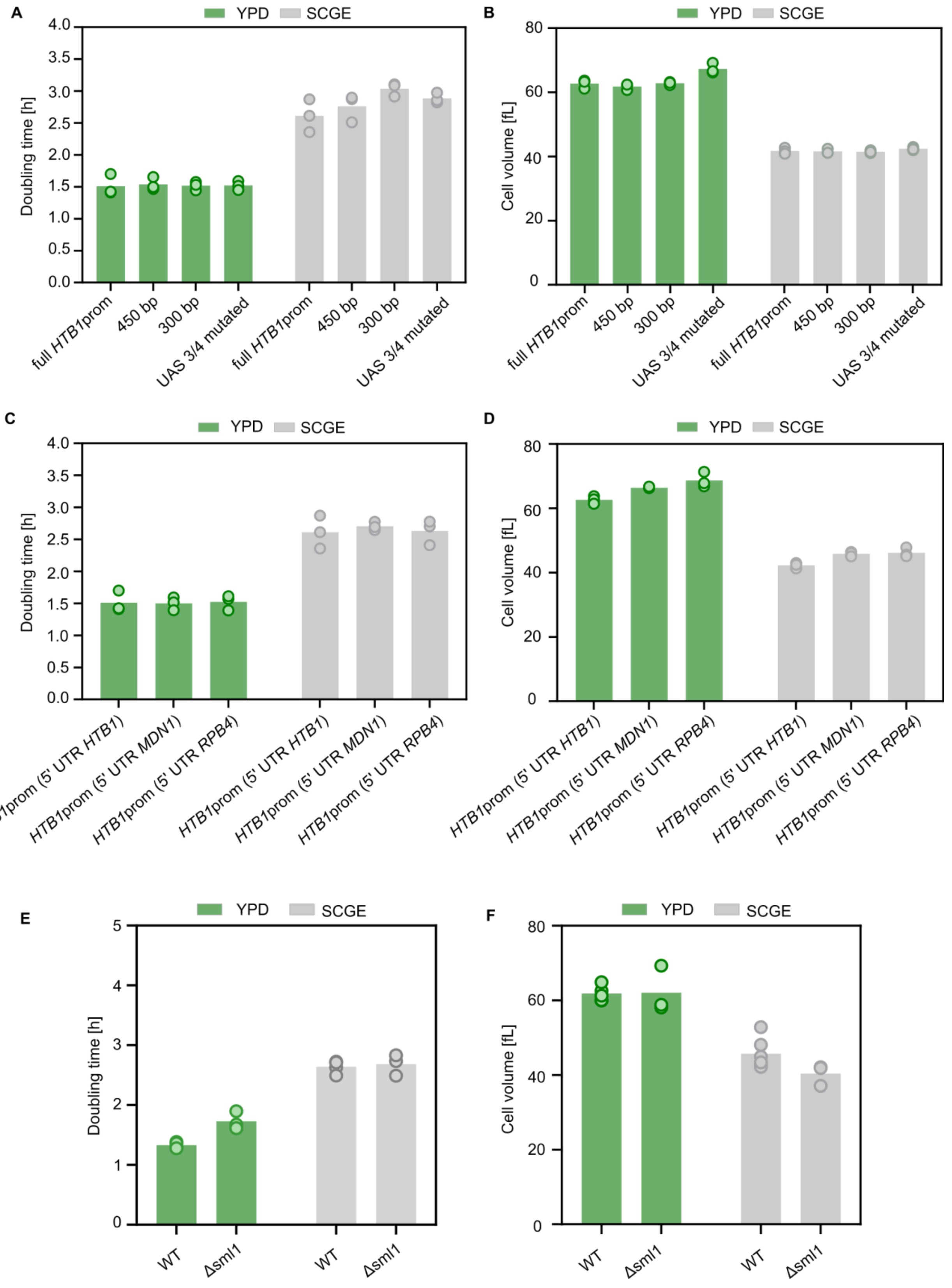
A
YPD
SCGE
Doubling time [h]
4.0
3.5
3.0
2.5
2.0
1.5
1.0
0.5
0.0
full HTB1prom
450 bp
300 bp
UAS 3/4 mutated
B
Cell volume [fL]
80
60
40
20
0
C
HTB1prom (5' UTR HTB1)
HTB1prom (5' UTR MDN1)
HTB1prom (5' UTR RPB4)
D
E
5
4
3
2
1
0
WT
Δsml1
F

**Supplementary figure 7. Growth phenotypes of strains used in figures 4, 5 and 6**. (**A**) Nutrient-specific population doubling times and (**B**) mean volumes of cells carrying an additional copy of the full-length, truncated or UAS3/UAS4 mutant *HTB1* promoter driving the expression of mCitrine. Bars represent the mean of n = 3 independent measurements shown as individual dots. (**C**) Nutrient-specific population doubling times and (**D**) mean volumes of cells expressing mCitrine with the *HTB1*, *MDN1* or *RPB4* 5' UTR. Bars represent the mean of n = 3 independent measurements shown as individual dots. (**E**) Population doubling times and (**F**) mean cell volumes of wildtype and *Δsml1* cells growing in YPD and SCGE. Bars represent the mean of n = 3-6 independent measurements shown as individual dots.

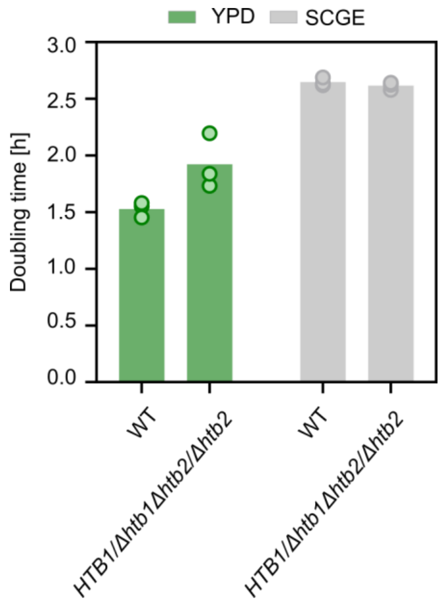


**Supplementary figure 8. Reduced histone amounts slow down cell growth in rich nutrients.** Doubling times calculated from growth curves of *HTB1/Δhtb1Δhtb2/Δhtb2* and wildtype diploid cells growing on fermentable and non-fermentable carbon sources. Bars represent the mean of n = 3 independent measurements shown as individual dots.

**Table 1.** Strains used in this study.

| Name | Genotype | Description | Origin | Figure |
|---|---|---|---|---|
| **ASY020-1** | *Mat α/a; ADE2/ADE2, URA3/ura3, leu2/LEU2* | Diploid wildtype strain | Anika Seel, Schmoller Lab | S8 |
| **CY14093** | *Mat α; sml1Δ::hphMX3* | Haploid *sml1Δ* strain | Christopher Bruhn | 6, S7 |
| **CY14098** | *Mat α; sml1Δ::hphMX3, rad53Δ::natMX6* | Haploid *sml1Δrad53Δ* strain | Christopher Bruhn | 6, S7 |
| **CY15164** | *Mat α; sml1Δ::hphMX3, rad53Δ::natMX6, spt21Δ::kanMX6* | Haploid *sml1Δrad53Δ spt21Δ* strain | Christopher Bruhn | 6, S7 |
| **DBY020-2** | *Mat α; ADE2, ura3::CglaTRP1-HTB1prom-mCitrine-ADH1term-URA3* | Haploid strain with additional copy of *HTB1* promoter driving mCitrine | This study | 4,5,S7 |
| **DBY021-3** | *Mat α; ADE2, ura3::CglaTRP1-HTB2prom-mCitrine-ADH1term-URA3* | Haploid strain with additional copy of *HTB2* promoter driving mCitrine | This study | 5 |
| **DBY054-8** | *Mat a/α ; ADE2/ADE2, leu2-3/LEU2, URA3/ura3-1, htb1Δ::CglaTRP1/ /HTB1, htb2Δ::HIS3 /htb2Δ::NatMX6* | Diploid *HTB1/htb1Δ, htb2Δ/htb2Δ* strain | This study | S8 |
| **DCY001-1** | *Mat α; ADE2, htb2::Htb2-linker-mCitrine-ADH1term-CglaTRP1, htb1::Htb1-linker-mCitrine-ADH1term-KlacURA3* | Haploid strain with *HTB1* and *HTB2* tagged with mCitrine | This study | 1, S2,S3 |
| **DCY002-2** | *Mat α; ADE2, ura3::CglaTRP1-HTB2prom-mCitrine-ADH11term-URA3,his3::ACT1prom-mKate2-ADH1term-HIS3* | Haploid strain with additional copy of *HTB2* promoter driving mCitrine and *ACT1* | This study | 3,S4 |

| | | | | |
|---|---|---|---|---|
| | | promoter driving mKate2 | | |
| **DCY006-1** | *Mat α ; ADE2, ura3::CglaTRP1-HTB1prom (mutated UAS3/UAS4)-mCitrine-ADH1term-URA3,his3::ACT1pr-mKate2-ADH1term-HIS3* | Haploid strain with additional copy of *HTB1* promoter, with mutated Spt10 binding sites in UAS3 and UAS4, driving mCitrine and *ACT1* promoter driving mKate2 | This study | 4,7 |
| **DCY008-8** | *Mat α; ADE2, ura3::CglaTRP1-HTB1prom-mCitrine-ADH1term-URA3,his3::ACT1prom-mKate2-ADH1term-HIS3* | Haploid strain with additional copy of *HTB1* promoter driving mCitrine and *ACT1* promoter driving mKate2 | This study | 3,S4 |
| **DCY011-1** | *Mat α; ADE2, ura3::CglaTRP1-HTB1prom- MDN1 5'UTR-mCitrine-ADH1term-URA3* | Haploid strain with additional copy of *HTB1* promoter expressing mCitrine with *MDN1* 5' UTR | This study | 5,S7 |
| **DCY012-1** | *Mat α; ADE2, ura3::CglaTRP1-HTB1prom- RPB4 5'UTR-mCitrine-ADH1term-URA3* | Haploid strain with additional copy of *HTB1* promoter expressing mCitrine with *RPB4* 5' UTR | This study | 5,S7 |
| **KCY021-1** | *Mat α; ADE2, ura3::CglaTRP1-300bpHTB1prom-mCitrine-ADH1term-URA3* | Haploid strain with additional 300 bp truncation of *HTB1* promoter driving mCitrine | Kora-Lee Claude, Schmoller Lab | 4,S7 |

| | | | | |
|---|---|---|---|---|
| **KCY022-1** | *Mat α; ADE2, ura3::CglaTRP1-450bpHTB1prom-mCitrine-ADH1term-URA3* | Haploid strain with additional 450 bp truncation of *HTB1* promoter driving mCitrine | Kora-Lee Claude, Schmoller Lab | 4,S7 |
| **KSY229-1** | *Mat α; ADE2, ura3::CglaTRP1-ACT1prom-mCitrine-ADH1term-URA3* | Haploid strain with additional copy of *ACT1* promoter driving mCitrine | This study | 3,S5 |
| **MMY116-2C** | *Mat α; ADE2* | Haploid wildtype strain | Skotheim lab stock | 1,2,3,5,S1,S3,S4,S6,S7 |

**Table 2.** Sequences of qPCR primers used in this study.

| Gene | qPCR primer direction | qPCR primer sequence (5‘-3‘) |
|---|---|---|
| **HTB1** | forward | TACACACATACAATGTCTGCTAAAG |
| | reverse | AGTGTCAGGGTGAGTTTGCTT |
| **HTB2** | forward | CCTCTGCCGCCGAAAAGAAA |
| | reverse | TCTTACCATCGACGGAGGTTG |
| **HTA1** | forward | GTTGCCAAAGAAGTCTGCCA |
| | reverse | CAGTTTAGTTCCTTCCGCCTT |
| **HTA2** | forward | TCGCCCAAGGTGGTGTTTT |
| | reverse | TGATTTGCTTTGTTTCTTTTCAACT |
| **HHF1** | forward | TACACCGAACACGCCAAGAG |
| | reverse | TTGCTTGTTGTTACCGTTTTCTT |
| **HHF2** | forward | ACGAAGAAGTCAGAGCCGTC |
| | reverse | ACCGATTGTTTAACCACCGATTG |
| **HHT1** | forward | CAATCTTCTGCCATCGGTGC |
| | reverse | ACTGATGACAATCAACAAACTATGA |
| **HHT2** | forward | AGCAAACACTCCACAATGGC |
| | reverse | CAAGGCAACAGTACCTGGCT |
| **ACT1** | forward | AGTTGCCCCAGAAGAACACC |
| | reverse | GGACAAAACGGCTTGGATGG |
| **MDN1** | forward | CATCAACAAACCTGACCAACTAATCC |
| | reverse | CATCAAGGTTTTCCAAAGTGGGC |
| **mCitrine** | forward | GAGCTGAAGGGCATCGACTT |
| | reverse | TTCTGCTTGTCGGCCATGAT |
| **RDN18** | forward | AACTCACCAGGTCCAGACACAATAAGG |
| | reverse | AAGGTCTCGTTCGTTATCGCAATTAAGC |

**Table 3.** Sequences of qPCR primers used for mRNA stability measurements shown in figure 2B.

| Gene | qPCR primer direction | qPCR primer sequence (5‘-3‘) |
|---|---|---|
| **HTB1** | forward | TGGCTGCGTATAACAAGAAGTCT |
| | reverse | CCAAAGGAAGTGATTTCATTATGC |
| **HTB2** | forward | TGCTCTATACTCAAACCAACAACA |
| | reverse | ATCTCTTCTTACCATCGACGGA |
| **ACT1** | forward | TATGTGTAAAGCCGGTTTTGC |
| | reverse | GACAATAC CGTGTTCAATTGGG |

**Table 4.** Promoter sequences used for the experiments shown in Fig. 5, in which *mCitrine* mRNA is expressed from the *HTB1 promoter* with *HTB1, MDN1* or *RPB4* 5' UTR.

| Gene feature | Position relative to start codon |
|---|---|
| ***HTB1* promoter** | - 817 bp to -127 bp relative to *HTB1* ORF |
| ***HTB1* 5' UTR** | 127 bp upstream of *HTB1* ORF |
| ***MDN1* 5' UTR** | 150 bp upstream of *MDN1* ORF |
| ***RPB4* 5' UTR** | 125 bp upstream of *RPB4* ORF |
